# A genome-scale phylogeny of Fungi; insights into early evolution, radiations, and the relationship between taxonomy and phylogeny

**DOI:** 10.1101/2020.08.23.262857

**Authors:** Yuanning Li, Jacob L. Steenwyk, Ying Chang, Yan Wang, Timothy Y. James, Jason E. Stajich, Joseph W. Spatafora, Marizeth Groenewald, Casey W. Dunn, Chris Todd Hittinger, Xing-Xing Shen, Antonis Rokas

## Abstract

Phylogenomic studies based on genome-scale amounts of data have greatly improved understanding of the tree of life. Despite their diversity, ecological significance, and biomedical and industrial importance, large-scale phylogenomic studies of Fungi are lacking. Furthermore, several evolutionary relationships among major fungal lineages remain controversial, especially those at the base of the fungal phylogeny. To begin filling these gaps and assess progress toward a genome-scale phylogeny of the entire fungal kingdom, we compiled a phylogenomic data matrix of 290 genes from the genomes of 1,644 fungal species that includes representatives from most major fungal lineages; we also compiled 11 additional data matrices by subsampling genes or taxa based on filtering criteria previously shown to improve phylogenomic inference. Analyses of these 12 data matrices using concatenation- and coalescent-based approaches yielded a robust phylogeny of the kingdom in which ∼85% of internal branches were congruent across data matrices and approaches used. We found support for several relationships that have been historically contentious (e.g., for the placement of Wallemiomycotina (Basidiomycota), as sister to Agaricomycotina), as well as evidence for polytomies likely stemming from episodes of ancient diversification (e.g., at the base of Basidiomycota). By examining the relative evolutionary divergence of taxonomic groups of equivalent rank, we found that fungal taxonomy is broadly aligned with genome sequence divergence, but also identified lineages, such as the subphylum Saccharomycotina, where current taxonomic circumscription does not fully account for their high levels of evolutionary divergence. Our results provide a robust phylogenomic framework to explore the tempo and mode of fungal evolution and directions for future fungal phylogenetic and taxonomic studies.

## Introduction

Fungi comprise an estimated 2-5 million species^1,2^, represent one of the most diverse and ancient branches of the tree of life, and play vital roles in terrestrial and aquatic ecosystems (Fig.1)^3^. Fungal organisms exhibit a wide variety of morphologies, developmental patterns, and ecologies, and are thought to have coevolved with plants through diverse modes of symbiosis, including parasitism, mutualism, and saprotrophy^4^. Many fungi are economically important as model organisms (e.g., the brewer’s yeast *Saccharomyces cerevisiae*, the fission yeast *Schizosaccharomyces pombe*, and the bread mold *Neurospora crassa*); food sources (e.g., mushrooms and truffles); cell factories for the production of diverse organic acids, proteins, and natural products (e.g., the mold *Aspergillus niger*); or major pathogens of plants and animals (e.g., the rice blast fungus *Magnaporthe grisea* and the amphibian chytrid fungus *Batrachochytrium dendrobatidis*), including humans (e.g., *Candida* species causing candidiasis, *Aspergillus* species causing aspergillosis, and *Cryptococcus* species causing cryptococcosis)^3,5,6^.

There are more than 200 orders of fungi that are classified into 12 phyla^6^ (for an alternative scheme of classification, see Ref.^7^). These 12 phyla are placed into six major groups: the subkingdoms Dikarya (which includes the phyla Ascomycota, Basidiomycota, and Entorrhizomycota) and Chytridiomyceta (which includes the phyla Chytridiomycota, Monoblepharidomycota, and Neocallimastigomycota), the phyla Mucoromycota, Zoopagomycota, and Blastocladiomycota, and the major group Opisthosporidia (which includes the phyla Aphelidiomycota, Cryptomycota/Rozellomycota, and Microsporidia, and is possibly paraphyletic)^6^. Evolutionary relationships among some of these groups, as well as among certain phyla and classes have been elusive, with morphological and molecular studies providing support for conflicting phylogenetic hypotheses or being equivocal in their support among alternatives^6,8^ (Supplementary Fig. 1).

**Figure 1.**
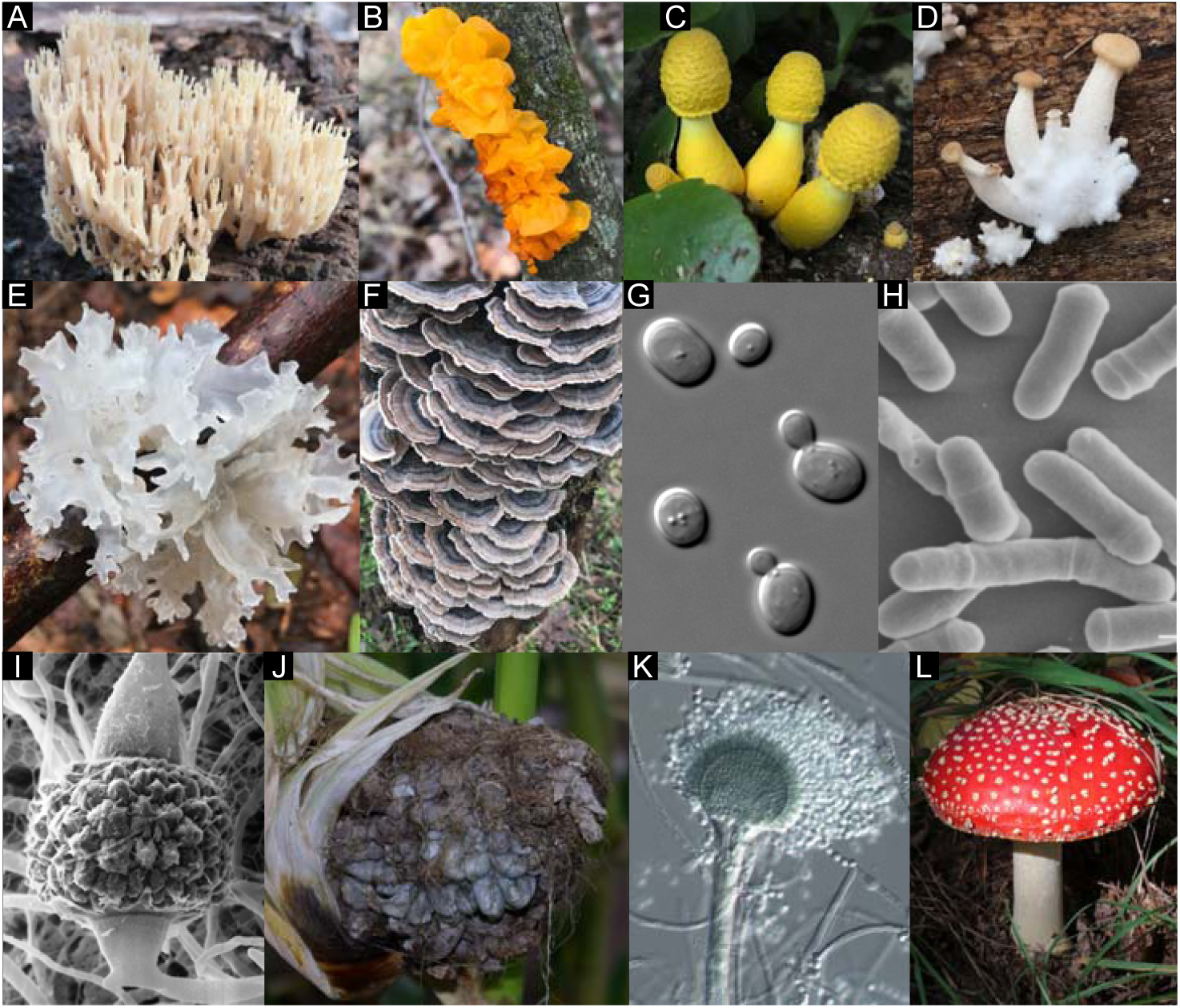
Diversity of major fungal lineages. Representative species for major fungal lineages. (A) Crown coral *Artomyces pyxidate* (Agaricomycotina, Basidiomycota). (B) Yellow brain fungus *Tremella mesenterica* (Pucciniomycotina, Basidiomycota). (C) Flowerpot parasol, *Leucocoprinus birnbaumii* (Agaricomycotina, Basidiomycota). (D) Pearl oyster mushroom, *Pleurotus ostreatus* (Agaricomycotina, Basidiomycota). (E) Snow fungus, *Tremella fuciformis* (Agaricomycotina, Basidiomycota). (F) Turkey tail, *Trametes versicolor* (Agaricomycotina, Basidiomycota). (G) Baker’s yeast *Saccharomyces cerevisiae* (Saccharomycotina, Ascomycota). (H) Fission yeast *Schizosaccharomyces pombe* (Taphrinomycotina, Ascomycota). (I) *Mucor mucedo* (Mucoromycotina, Mucoromycota). (J) Corn smut *Ustilago maydis* (Ustilaginomycotina, Basidiomycota). (K) *Penicillium digitatum* (Pezizomycotina, Ascomycota). (L) Fly agaric *Amanita muscaria* (Agaricomycotina, Basidiomycota). A-F, Photograph courtesy of Jacob L. Steenwyk. G, J, L, Images are available to the public domain through https://commons.wikimedia.org/wiki/. H, Photograph reproduced with permission of David O. Morgan. I, Photograph courtesy of Kerry O’Donnell and Jason Stajich.

Several notable cases of phylogenetic ambiguity exist. For example, relationships among the three phyla that comprise Opisthosporidia are ambiguous, especially the placement of Aphelidiomycota (Supplementary Fig. 1). This is likely due to the parasitic lifestyles and highly reduced morphologies of many of the organisms involved that coincide with highly reduced and very rapidly evolving genomes (e.g., Microsporidia), that render their evolutionary placement challenging^9,10^. Ambiguity also exists with respect to the placement of Blastocladiomycota – either as a sister to the rest of fungi excluding Opisthosporidia or as a sister to the rest of fungi excluding Opisthosporidia and Chytridiomyceta^6,11–13^.

Although aquatic, Blastocladiomycota have several traits found in terrestrial fungi, making their placement on the fungal phylogeny key for understanding the evolution of diverse fungal traits^6,14^. Mucoromycota and Zoopagomycota were previously classified into the phylum Zygomycota based on the production of coenocytic hyphae and sexual reproduction by zygospores. Mucoromycota and Zoopagomycota were previously classified into the phylum Zygomycota based on the production of coenocytic hyphae and sexual reproduction by zygospores. However, after the arbuscular mycorrhizal fungi were segregated from Zygomycota into a new phylum (Glomeromycota)^15^, the Zygomycota became paraphyletic^8,14^ and was abandoned in favor of a classification of the two major lineages (Mucoromycota and Zoopagomycota)^8^. Other classifications have retained Glomeromycota and seven additional phyla for the former Zygomycota^7^. Finally, evolutionary relationships of major lineages within Chytridiomyceta (i.e., between Chytridiomycota, Neocallimastigomycota, Monoblepharidomycota), Basidiomycota (i.e., among Agaricomycotina, Pucciniomycotina, and Ustilaginomycotina), and phylum Ascomycota (e.g., between classes in Taphrinomycotina) are also elusive^6,8^ (Supplementary Fig. 1).

In retrospect, previous molecular phylogenetic analyses have relied primarily on a handful of loci from many taxa that often provided little resolution of the deep internal branches (e.g., 6 genes / 199 taxa^16^), or genomic data with relatively scarce taxon sampling (e.g., 53 genes / 121 taxa^17^; 192 genes / 46 taxa^18^; 650 genes / 104 taxa^19^; 455 genes / 72 taxa^20^). However, phylogenomic studies of specific fungal lineages that are well sampled, such as Saccharomycotina (e.g., 2,408 genes / 332 taxa^21^) and Ascomycota (e.g., 815 genes / 1,107 taxa^22^), suggest that denser gene and taxon sampling hold great potential for resolving relationships that previously seemed intractable. For example, in a phylogeny of budding yeasts (subphylum Saccharomycotina, phylum Ascomycota) inferred from analyses of 1,233-gene, 86-taxon data matrix^23^, we noticed that the placement of the family Ascoideaceae, which was represented by the genome of *Ascoidea rubescens*, was sensitive to the inclusion of a single gene with unusually strong phylogenetic signal^24^. In contrast, analysis of a 2,408-gene, 332-taxon data matrix that included four taxa from Ascoideaceae robustly resolved the placement of this lineage on the tree of budding yeasts and ameliorated the gene’s undue impact^21^.

A robust phylogenetic framework for Fungi based on a broad sampling of genes and taxa is key for understanding the evolution of the group and would greatly facilitate larger-scale studies in fungal comparative biology, ecology, and genomics. In recent years, the 1000 Fungal Genomes Project (https://mycocosm.jgi.doe.gov/programs/fungi/1000fungalgenomes.jsf) has greatly expanded the availability of genomes from diverse understudied taxa across the fungal tree of life^25^. Additionally, efforts focused on specific ecological or evolutionary groups, such as the Y1000+ Project (http://y1000plus.org) that aims to sequence all known species of the subphylum Saccharomycotina^26^, the Dothideomycetes project that aims to study plant pathogenic fungi^27^, and the *Aspergillus* genome project that aims to examine the metabolic dexterity of this diverse genus of fungi^28^ (https://jgi.doe.gov/aspergillus-all-in-the-family-focused-genomic-comparisons), have greatly increased the availability of genomes from specific lineages.

The availability of genomic data from a substantially expanded and more representative set of fungal species offers an opportunity to reconstruct a genome-scale fungal tree of life and examine its support for relationships that have heretofore remained contentious or poorly resolved (Supplementary Fig. 1). To this end, we analyzed data from 1,644 available fungal genomes that include representatives from most major lineages and provided a robust phylogenomic framework to explore the evolution of the fungal kingdom.

## Results and Discussion

### A pan-fungal phylogenomic matrix with high taxon sampling and occupancy

To assemble a phylogenomic data matrix, we sampled 1,707 publicly available genomes from NCBI (one representative genome from every species; retrieved on January 30, 2020), representing every major lineage across fungi (1,679 taxa) and selected outgroups (28 taxa) based on the current understanding of the Opisthokonta phylogeny^29,30^; the sole exceptions were the Aphelidiomycota and Entorrhizomycota phyla, for which no genomes were available as of January 30, 2020 (Supplementary Table 1).

To filter out low quality genomes, we analyzed each genome using BUSCO^31^ with the Fungi OrthoDB v9 database^32^, which contains 290 genes. This analysis resulted in the removal of the genomes of 35 fungal species, each of which had fewer than 100 single-copy BUSCO genes. The average genome assembly completeness for the remaining 1,672 taxa was 92.32% (average of 267.74 / 290 BUSCO genes). The full data matrix contains 124,700 amino acid sites from 290 BUSCO genes (90.6% taxon-occupancy per BUSCO gene, an average length of 430 residues per gene after trimming, and missing data of 15.64% (84.36% matrix occupancy)) across 1,672 taxa (1,644 fungal taxa and 28 outgroups). Annotations and characteristics of each BUSCO gene, including its length and taxon occupancy, are presented in Tables S1 and S2.

To conduct sensitivity analyses for potential systematic errors or biases that may influence the accuracy of phylogenetic inference, we generated 11 additional data matrices by subsampling genes (8 data matrices) or taxa (3 data matrices) from the full data matrix. The examined biases include the removal of genes (e.g., based on shorter alignment length and higher evolutionary rate) or taxa (e.g., based on LB-score, or by removing rogue taxa) according to filtering criteria previously shown to improve phylogenomic inference (Supplementary Fig. 2; also see Methods for detailed information for each matrix)^33,34^.

**Figure 2.**
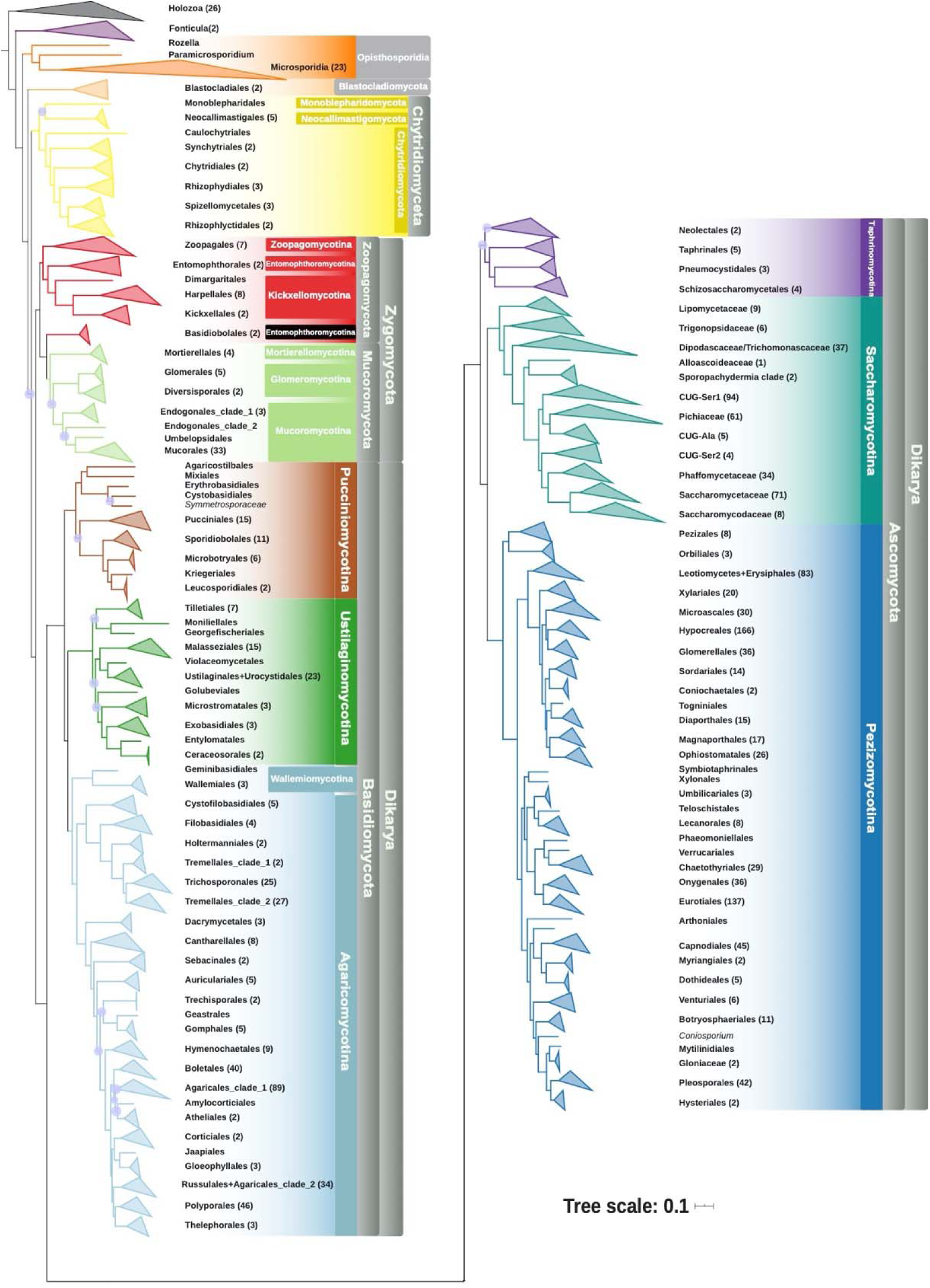
Genome-scale phylogeny of 1,644 species spanning the diversity of Fungi. The topology shown is derived from maximum likelihood analysis using a concatenation single-model approach on the full data matrix (1,672 taxa (1,644 fungi and 28 outgroups) and 290 genes; *lnL* = -78287339.984). Internal branches supported with 100% bootstrap values are not shown; those with values lower than 100% are denoted by purple dots. Each tip corresponds to the order-level ranking derived from NCBI taxonomy (except for subphylum Saccharomycotina, where each tip corresponds to each one of the 12 major clades to reflect the current understanding of Saccharomycotina phylogeny^21^).

### A robust phylogenetic framework to explore fungal evolution

To infer the fungal phylogeny, we used concatenation-based single model (unpartitioned), concatenation-based data-partitioning (one partition per gene), and coalescent-based approaches on the full data matrix as well as on the 11 additional data matrices (Supplementary Fig. 2). The gene occupancy for every taxon in each data matrix is shown in Supplementary Table 2. These analyses produced 33 phylogenetic trees: 12 from concatenation-based single model analyses, nine from concatenation-based data-partitioning analyses (phylogenies were not inferred from three matrices for reasons of computational efficiency), and 12 from coalescent-based analyses; see methods for more details. We found that ∼85% (1,414 / 1,669 of bipartitions (or internodes / internal branches) were recovered consistently across these 33 phylogenies, suggesting that a large fraction of internal branches of the fungal phylogeny were robustly supported (Supplementary Fig. 4).

Notable examples of relationships that were recovered in all 33 phylogenies included the placement of the cellular slime mold *Fonticula* as sister to fungi and the placement of Opisthosporidia as sister to the rest of fungi (Figs. 2, 3)^30,35^. Our analyses also robustly placed Wallemiomycotina within Basidiomycota, which has historically been contentious since it has been placed sister to^36^, within^37^, or outside^38^ of Agaricomycotina. All of our analyses placed Wallemiomycotina (which contains both Geminibasidiales and Wallemiales) as sister to Agaricomycotina with strong support (BS = 100; LPP = 100) (Figs. 2 and 3).

**Figure 3.**
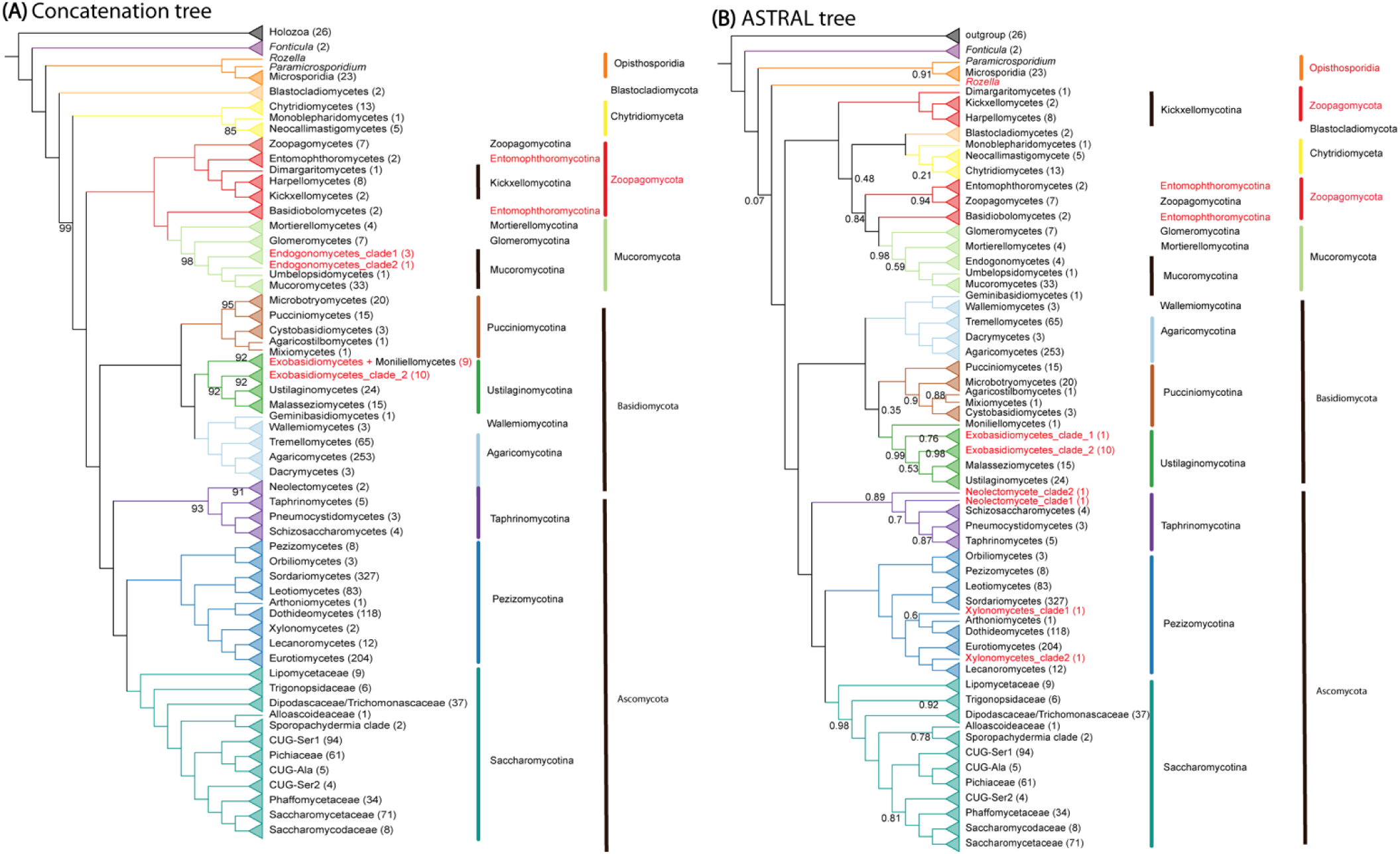
Incongruence between concatenation- and coalescent-based phylogenies of Fungi. Topologies derived from maximum likelihood analysis using (A) a concatenation single model (LG+G4) approach and (B) a coalescent-based approach. Support for internal branches with 100% bootstrap values (for A) or posterior probabilities of 1.0 (for B) is not shown. Each tip corresponds to the class-level ranking derived from NCBI taxonomy (except for subphylum Saccharomycotina, where each tip corresponds to each one of the 12 major clades to reflect the current understanding of Saccharomycotina phylogeny^21^). Taxa in red correspond to groups inferred to be paraphyletic by the topology shown.

**Figure 4.**
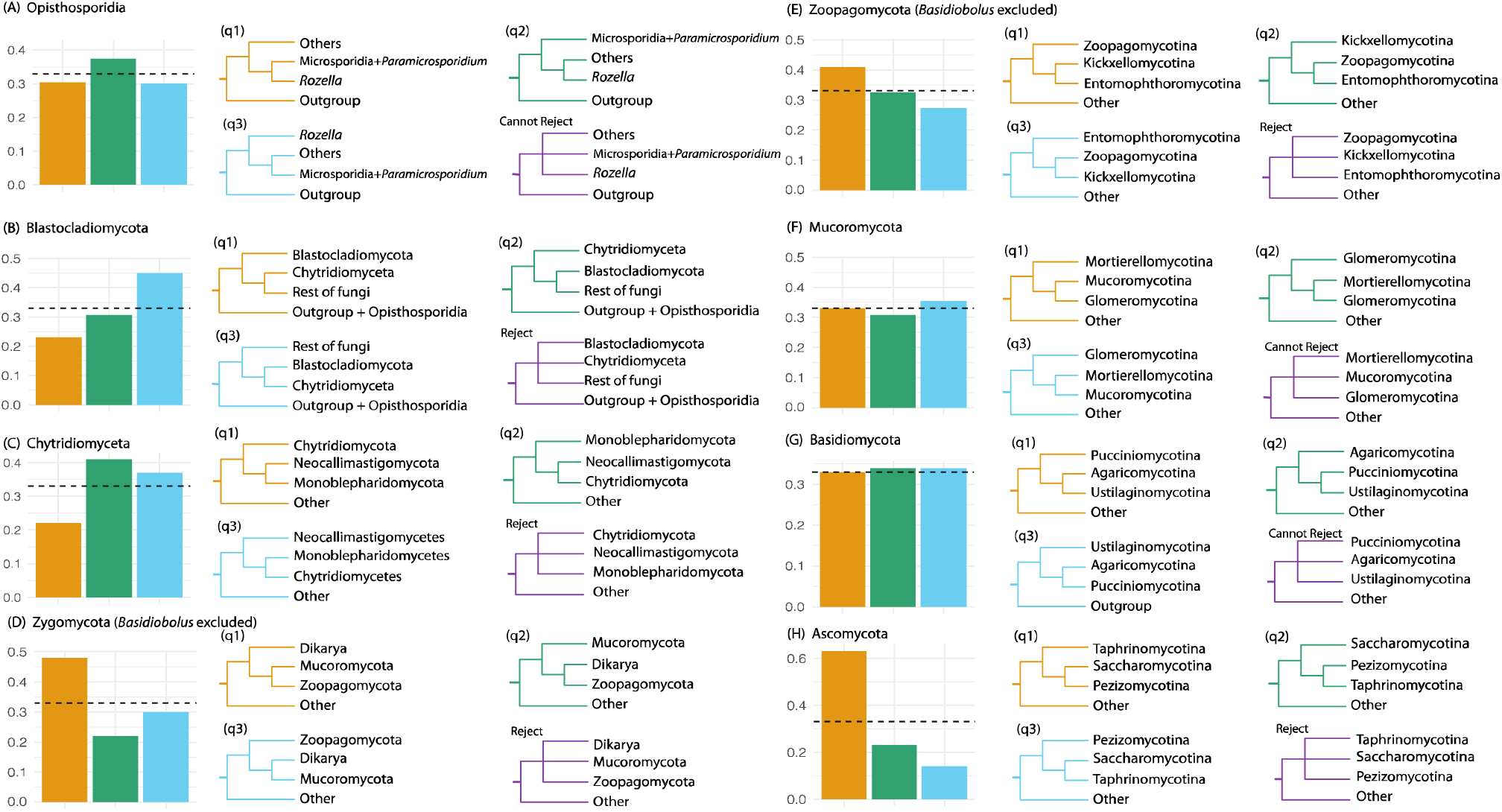
Examination of support among individual gene trees for alternative hypotheses for contentious relationships in the fungal phylogeny. The gene-tree quartet frequencies (bar graphs) for alternative branching orders for contentious relationships in the fungal phylogeny. (**A**) Opisthosporidia diversification. (**B**) Blastocladiomycota and Chytridiomyceta diversification. (**C**) Chytridiomyceta diversification. (**D**) Monophyly of Zygomycota. (**E**) Zoopagomycota diversification. (**F**) Mucoromycota diversification. (**G**) Basidiomycota diversification. (**H**) Ascomycota diversification. Orange bars and topologies reflect the relationships inferred using a concatenation-based single model approach on the full data matrix; blue and green bars and trees correspond to the two alternative hypotheses (supported by the two alternative resolutions of each quartet). The purple tree shows whether a polytomy scenario can be rejected by the quartet analysis or not. Dashed horizontal lines mark expectation for a hard polytomy.

In general, robustly supported relationships were more commonly found in parts of the tree with better taxon sampling. For Ascomycota, the phylum with the best sampling of taxa in our data matrix, ∼94% of bipartitions (1,036 / 1,101) were consistently recovered across the 33 phylogenies. For example, we found that all 33 phylogenies strongly supported Taphrinomycotina as the sister lineage to a clade of Saccharomycotina and Pezizomycotina (BS = 100; LPP = 100; q1 = 0.62) (Figs. 3, 4H). Similarly, all phylogenies strongly supported a clade consisting of Pezizomycetes and Orbiliomycetes as the sister group to the remaining Pezizomycotina (Fig. 3). Both Saccharomycotina (332 taxa with representatives of all 12 major clades included) and Pezizomycotina (761 taxa with 9 of the 17 known classes included) are the most well-sampled major lineages in our data matrix (Supplementary Table 2), suggesting that a combination of increased taxon sampling and genome-scale amounts of data can improve the resolution of the fungal tree of life. Importantly, relationships among the 12 major clades of the subphylum Saccharomycotina and relationships among higher taxonomic ranks within Ascomycota recovered by our analyses are essentially the same as those of previous studies performed using different sets of genes and taxa^21,22^.

Finally, we note that a recent study used the alignment-free Feature Frequency Profile (FFP) method to reconstruct a broad sketch of the fungal tree of life based on proteome data from over 400 fungal genomes^39^. However, our recent simulation study showed that the performance of the FFP method is much worse than concatenation- and coalescent-based approaches for reconstructing the phylogeny of major and ancient lineages^40^, such as fungi. The poor performance of the FFP method explains why many relationships reported by Choi and Kim^39^, such as the sister group relationships for Chytridiomyceta + Zygomycota and for Taphrinomycotina + Saccharomycotina, strongly contradict the current consensus view of the fungal tree of life^6,22^.

### Most instances of incongruence stem from differences between concatenation- and coalescent-based phylogenies

By examining the distribution of incongruence across the 33 phylogenies, we found that the 21 phylogenies obtained from concatenation-based single model and data-partitioning analyses were largely congruent; an average of 98.6% (1,645 /1,669) of bipartitions were recovered consistently (Supplementary Fig. 4). The high similarity between these two approaches is consistent with findings of previous phylogenomic studies^21^.

In contrast, from a total of 255 incongruent bipartitions found across the 33 phylogenies, 145 (average = 58.9%) were mainly due to whether the data matrix was analyzed by concatenation or coalescence (Supplementary Fig. 4). Furthermore, these incongruent bipartitions were more concentrated in branches toward the base of the fungal phylogeny (Fig. 3). By examining incongruence at the taxonomic levels of order, class, and phylum, we found four ranks that were recovered as non-monophyletic in concatenation-based analyses compared to seven non-monophyletic ranks in coalescent-based analyses (Fig. 3, Supplementary Table 3). Coalescent-based trees contradict well-established relationships supported by most previous phylogenetic studies, as well as by our concatenation-based analyses, such as the sister group relationship of Rozellomycota and Microsporidia^35,41^ and the monophyly of Zoopagomycota (excluding *Basidiobolus*)^18^ (Fig. 3B).

The observed differences between concatenation-based and coalescent-based analyses may stem from the fact that a substantial number of internodes in individual gene trees are not well supported. We found an average of 4.99%, 6.69%, 10.74%, and 19.18% of internodes in individual gene trees that received ultrafast bootstrap support values lower than 33%, 50%, 75%, and 95%, respectively. Given that values above 95% are considered as strong support^42^, these results suggest that nearly one in five internodes in individual gene trees lacks robust support. Since our ASTRAL-based analyses use directly these gene trees to infer the coalescent-based species trees, their accuracy may be disproportionally be affected (compared to the concatenation-based species trees) by the poor resolution of individual gene trees.

An alternative, not mutually exclusive, explanation may be that there are, on average, 430 amino acid sites in each gene alignment, whereas the average number of internodes in each gene tree is 1,501. Thus, there may be more free parameters in individual gene tree estimates (each branch is considered a free parameter in maximum likelihood) than there are data points, which may affect gene tree accuracy and support due to over-parameterization.

Another possible explanation is that 290 genes are not sufficient to robustly resolve all internal branches of a tree with hundreds of taxa. The number of genes in a phylogenomic data matrix is known to impact the accuracy of both concatenation-based ^43^ and coalescent-based inference^44^. Moreover, the taxon-occupancy values for non-Dikarya fungi (average of 207.02 / 290 BUSCO genes; 71.39%) are lower than the ones of Dikarya (average of 279.59 / 290 BUSCO genes; 96.41%). Consequently, the placements of non-Dikarya taxa are based on many fewer genes and gene trees.

Notwithstanding the debate on which of the two approaches is better or more appropriate for estimating species phylogenies^45,46^, these results suggest that concatenation-based phylogenies of this phylogenomic data matrix are likely more trustworthy than coalescent-based phylogenies due to the poor resolution of individual gene trees.

### Incongruence among major lineages and identification of ancient radiations

Although ∼85% of internodes in our phylogeny of Fungi were robustly supported irrespective of approach and data matrix used, the remaining ∼15% showed incongruence between analyses. Below, we discuss certain key incongruent relationships of interest. For each case, we present the results from our concatenation- and coalescent-based analyses, and place our results in the context of the published literature. For key contentious branches of the fungal phylogeny, we also tested whether the data from the 290 gene trees rejected the hypothesis that the branch in question represents a polytomy (Fig. 4). Briefly, the polytomy test evaluates whether the frequencies of quartet trees (obtained from all the gene trees) are significantly different for a branch of interest^47^. For every quartet tree, there are three possible topologies (i.e., three alternative hypotheses noted as q1, q2, q3) of how the taxa are related. The test measures the frequencies of the quartet trees present in all gene trees; if there are no significant differences in their frequencies, then the hypothesis that the branch in question is a polytomy cannot be rejected. Given that the quartet frequencies are obtained from the individual gene trees, the analyses of Fig. 4 generally reflect the results of the coalescent-based analyses.

#### Opisthosporidia

Opisthosporidia are a group of reduced, endoparasite taxa that includes Rozellomycota, Microsporidia (parasites of animals), and Aphelidiomycota (parasites of algae for which no genomes are currently available) (Supplementary Fig. 1). Within Opisthosporidia, our concatenation-based analyses strongly supported a clade of Rozellomycota and Microsporidia (Figs. 2 and 3A). To date, only two genomes from the Rozellomycota have been sequenced, namely *Paramicrosporidium saccamoebae*^*35*^ and *Rozella allomycis*^*41*^. Both concatenation and coalescent-based analyses placed *P. saccamoebae* as sister to Microsporidia, suggesting that Rozellomycota is paraphyletic (Figs. 2 and 3). These results are largely consistent with previous gene content and phylogenetic analyses that *P. saccamoebae* is more closely related to Microsporidia than to other Rozellomycota^35^ (Supplementary Fig. 1). In contrast, the two approaches differed in the placement of *R. allomycis* (Fig. 3). Whereas concatenation-based analyses placed *R. allomycis* sister to the *P. saccamoebae* + Microsporidia clade (Fig. 3A), coalescent-based analyses placed *R. allomycis* as sister to the remaining non-Opisthosporidia fungi with very low support (LPP = 0.07) (Fig. 3B). Finally, quartet tree support for the concatenation-based placement (q1 = 0.31) was lower than the coalescent-based placement (q2 = 0.38) but a polytomy scenario could not be rejected (Fig. 4A).

Given that only two genomes from Rozellomycota and none from Aphelidiomycota are available, the lack of resolution within Opisthosporidia may be due to scarce taxon sampling. Although previous phylogenomic analyses based on a single transcriptome from Aphelidiomycota placed this phylum as sister to free-living fungi^9^, which would render Opisthosporidia paraphyletic, further studies with more taxa will be necessary to confidently resolve relationships in this lineage.

#### Blastocladiomycota, Chytridiomyceta, and the rest of fungi

Within zoosporic fungi, the relationship between Blastocladiomycota and Chytridiomyceta, and the rest of fungi (excluding Opisthosporidia) remains ambiguous^6,11–13^. Our concatenation analyses placed Blastocladiomycota as sister to a clade of Chytridiomyceta and the rest of fungi with strong support (BS = 99) (Fig. 3A). In contrast, coalescent-based analyses supported a sister taxon relationship between Blastocladiomycota and Chytridiomyceta with strong support (LPP = 1.00) (Fig. 3B). The quartet-based analyses showed low support for the concatenation-based placement (q1 = 0.24), intermediate support for Chytridiomyceta as sister to a clade of Blastocladiomycota and the rest of fungi (q2 = 0.31), and strong support for the coalescent-based placement (q3 = 0.45) (Fig. 4B). The low resolution of relationships between Blastocladiomycota and Chytridiomyceta in our coalescent-based analysis might be due to the lower taxon occupancy in these two clades (average of taxon occupancy: 73.68% in Chytridiomyceta; 42.59% in Blastocladiomycota) (Supplementary Table 2). Blastocladiomycota have characteristics that more resemble terrestrial fungi, such as well-developed hyphae, closed mitosis, cell walls with β-1-3-glucan, and a Spitzenkörper^48,49^. Thus, understanding the true branching order has important implications for the evolution of key traits and processes (e.g., life cycles, mitosis)^6^.

Within the subkingdom Chytridiomyceta, the phylogenetic relationships among the Monoblepharidomycota, Chytridiomycota, and Neocallimastigomycota are also uncertain^12,50^. Our concatenation analyses recovered Chytridiomycota as the sister group to a clade of Monoblepharidomycota and Neocallimastigomycota (BS = 85) (Fig. 3A), whereas coalescent analyses recovered Monoblepharidomycota as the sister to Chytridiomycota and Neocallimastigomycota (LPP = 0.18) (Fig. 3B). The quartet-based analyses showed lower support for the concatenation-based placement (q1 = 0.22) than for the coalescent-based placement (q2 = 0.41) or the third alternative hypothesis (q3 = 0.38) (Fig. 4C). Given that one genome was sampled from Monoblepharidomycota, 13 genomes were sampled from Chytridiomycota, and five genomes were sampled from Neocallimastigomycota, additional sampling of taxa, and perhaps genes as well, will be necessary for the confident resolution of relationships within Chytridiomyceta.

#### Monophyly of Zoopagomycota and Mucoromycota

The monophyly of Zygomycota was not supported in recent phylogenetic studies^12,16,18,50^. Because of the uncertain relationships among these fungi, there have been several classifications that split them into multiple subphyla and phyla^14,18^. Our concatenation analyses strongly supported the monophyly of Zoopagomycota and Mucoromycota, previously known as Zygomycota (BS = 100) (Fig. 3A). Coalescent analyses recovered the monophyly Mucoromycota, although as mentioned earlier, Chytridiomyceta and Blastocladiomycota are nested within Zoopagomycota in these coalescent-based phylogenies (Fig. 3B). The quartet-based analysis shows that the quartets for the monophyly of Zoopagomycota and Mucoromycota received the highest support (q1=0.48; Fig. 4D).

However, we found one data matrix that recovered the paraphyly of Zygomycota, albeit with very low support (BS = 28), stemming from the Top100_slow-evolving data matrix (Supplementary Fig. 6). This recovered topology is largely consistent with previous analyses and Zoopagomycota is also recovered as monophyletic (BS = 28).

To further explore the effect of gene sampling on the resolution of Zygomycota in different phylogenomic data matrices, we next quantified the support of phylogenetic signal over two alternative hypotheses (T1: Zygomycota-monophyly; T2: Zygomycota-paraphyly) using our Subset_Dikarya data matrix (see Methods) and a previously published 192-gene, 46-taxon data matrix (Spatafora2016_46taxa_192genes data matrix)^51^ (Supplementary Fig. 7). By calculating gene-wise log-likelihood scores between T1 and T2 (ΔlnL) for every gene in each matrix, we found that the proportions of genes supporting T1 versus T2 were similar in both data matrices (95 of 192: 49.5% vs 97 of 192: 50.5% in the Spatafora2016_46taxa_192genes matrix; 161 of 290: 55.5% vs 129 of 290: 44.5% in the Subset_Dikarya data matrix) (Supplementary Fig. 7), even though the results of our study support Zygomycota monophyly^52^ and those of other studies support Zygomycota paraphyly^12,16,18^. Thus, phylogenomic analyses of Zygomycota should be interpreted with caution until further taxon and gene sampling of taxa from the lineages in question sheds more light into this part of the fungal phylogeny.

#### Zoopagomycota paraphyly

We found that Zoopagomycota was paraphyletic because two *Basidiobolus* species were placed as the sister group to Mucoromycota (Figs. 2 and 3). The phylogenetic placement of *Basidiobolus* in previous phylogenetic analyses based on genomic^18^ or multigene^18,53^ studies was unstable, and a recent study has suggested that many genes in *Basidiobolus* genomes might have been acquired from Bacteria through horizontal gene transfers^54^. Notably, removal of the two *Basidiobolus* taxa in the removal-of-rogue-taxa data matrix did not alter the monophyly of Zygomycota (Supplementary Fig. 5), suggesting that this result was not affected by the topological instability of *Basidiobolus*.

**Figure 5.**
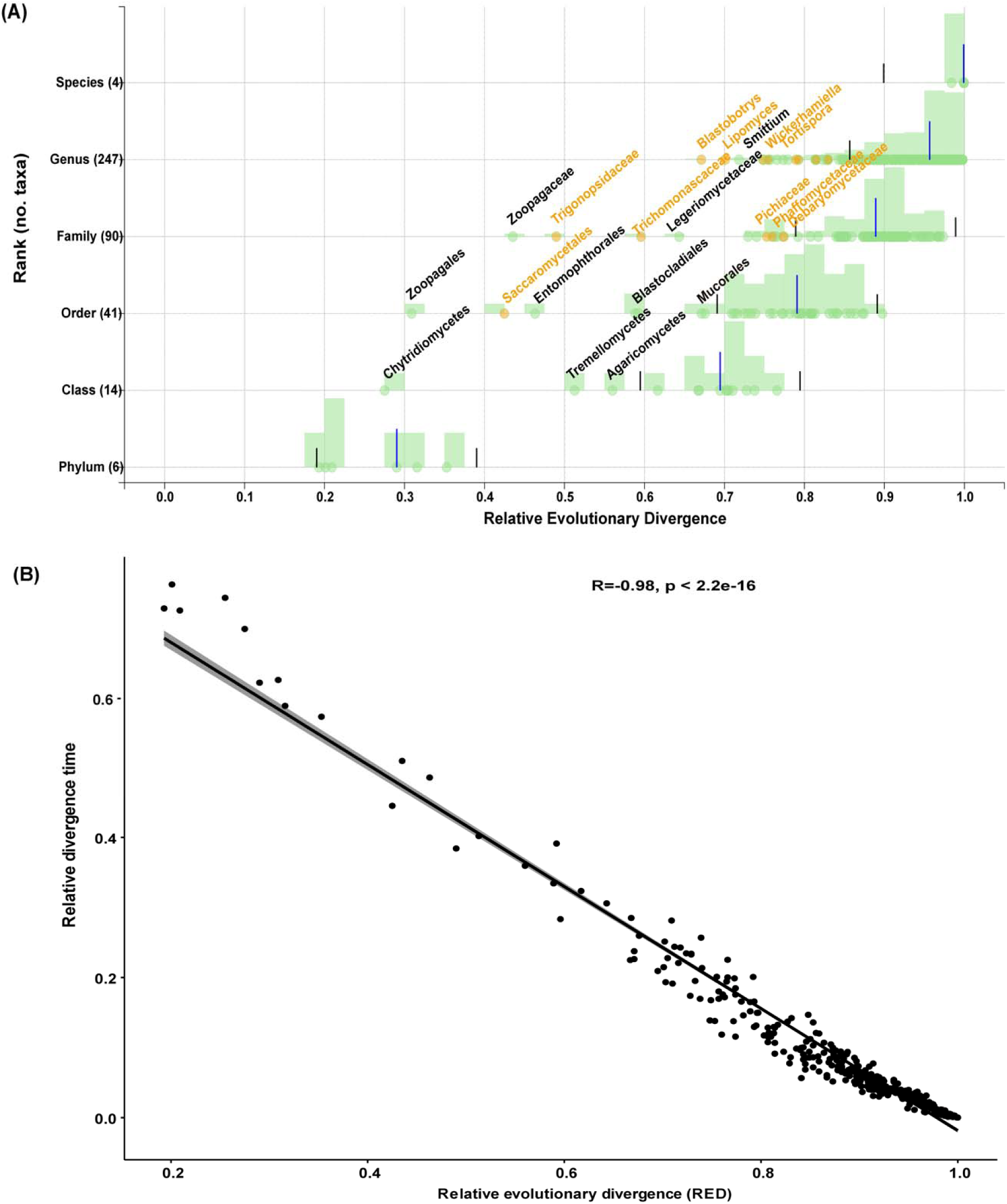
Higher level taxonomic ranks generally reflect levels of evolutionary divergence across the fungal kingdom. (**A**) Relative evolutionary divergence (RED) of taxa defined by the NCBI taxonomy based on the topology inferred from the concatenation-based single model approach. Each data point (green or orange circle) represents a taxon distributed according to its RED value (x-axis) and its taxonomic rank (y-axis). Blue bars correspond to median RED values and black bars to the RED intervals (+/- 0.1) for each rank. Orange circles represent taxa belonging to the subphylum Saccharomycotina (Ascomycota), which are the most notable instance of an underclassified lineage in the fungal kingdom. Note that the RED approach does not assign values for ranks with a single subordinate rank (e.g., class Saccharomycetes contains a single order, Saccharomycetales; thus, a RED value has only been assigned for Saccharomycetales and only the order’s value is plotted on the graph). Only a subset of taxon names is shown here; results for all taxa are reported in Supplementary Table 4. (**B**) The Pearson correlation coefficient (Pearson’s r) between the RED values and relative divergence time estimated using relaxed-molecular clock approaches for all internal nodes.

#### Major relationships within Zoopagomycota

Within Zoopagomycota, the monophyly of Zoopagomycotina, Kickxellomycotina, and Entomophthoromycotina (without *Basidiobolus*) was well supported in concatenation-based analyses, with Zoopagomycotina as sister to Kickxellomycotina and Entomophthoromycotina with strong support (BS = 100) (Fig. 2). This relationship is also supported in our quartet-based analysis (q1 = 0.41; q2 = 0.32; q3 = 0.27) (Fig. 4E). In contrast, as mentioned previously, coalescence failed to recover a monophyletic Zoopagomycota (without *Basidiobolus*; Fig. 3B).

#### Relationships within Mucoromycota

Within Mucoromycota, the concatenation-based analysis moderately supported Mortierellomycotina as sister to Mucoromycotina and Glomeromycotina (BS = 98), whereas the coalescent-based analysis placed Glomeromycotina sister to the remaining Mucoromycota with low support (LPP = 0.61) (Fig. 3). Quartet-tree support for the concatenation-based phylogeny was largely similar to the two alternative hypotheses (q1 = 0.33; q2 = 0.31; q3 = 0.36) (Fig. 4F), suggesting that a polytomy best explains relationships between subphyla of Mucoromycota based on current evidence. Nevertheless, the small number of genomes sampled suggests that these inferences may be subject to revision.

#### Evidence for a polytomy at the base of Basidiomycota

Subkingdom Dikarya comprises most of the described fungal diversity (∼97%), comprising Ascomycota, Basidiomycota, and Entorrhizomycota (for which no genome is currently available). Even though Basidiomycota have much denser taxon sampling than most other fungal lineages, reconstruction of the relationships among Pucciniomycotina, Ustilaginomycotina, and Agaricomycotina + Wallemiomycotina has proven challenging^37,39,55,56^. We too found discordant topologies between concatenation- and coalescent-based analyses (Fig. 3) and nearly equal support for the three alternative hypotheses (Fig. 4G). Concatenation analyses placed Ustilaginomycotina with Agaricomycotina + Wallemiomycotina (BS = 100), whereas coalescence ones supported Pucciniomycotina with Ustilaginomycotina (LPP = 0.41). Notably, we found that gene-tree quartet support for the three alternative hypotheses was consistent with a polytomy (q1 = 0.33, q2 = 0.34, q3 = 0.34) (Fig. 4G), suggesting that the origin of major lineages within Basidiomycota may be the result of an ancient diversification. Importantly, a previous study suggested that the divergence of Basidiomycota occurred after the diversification of extant embryophytes (481 - 452 million years ago)^4^, thus the ancient diversification of these major Basidiomycota lineages might have been driven by their association with early terrestrial embryophytes. For example, most Agaricomycotina are decomposers of mycorrhizae, whereas most Pucciniomycotina are obligate pathogens to all main lineages of embryophytes and nearly all Ustilaginomycotina are parasites specific to angiosperms^4^.

### Higher-level taxonomic ranks generally reflect levels of evolutionary divergence across the fungal kingdom

To evaluate whether higher taxonomic ranks of fungi exhibit comparable levels of evolutionary divergence, we normalized the fungal taxonomy ranks retrieved from the National Center for Biotechnology Information using the relative evolutionary divergence (RED) approach^57^. The RED approach normalizes the inferred phylogenetic distances between the last common ancestor of fungi (RED = 0) to all extant fungal taxa (RED = 1) to provide an approximation of the relative time of divergence (Fig. 5A). We note that the values obtained through application of the RED approach to the fungal phylogeny are broadly consistent to the relative divergence time estimated using relaxed-molecular clock approaches (Pearson’s correlation coefficient *r* = -0.98, *P*-value < 2.2e-16) (Fig. 5B). Note that the RED approach does not assign values for ranks with a single subordinate rank (e.g., class Saccharomycetes contains a single order, Saccharomycetales; thus, the RED value will be only assigned to the order Saccharomycetales (Fig. 5A)); consequently, the number of taxonomic ranks under evaluation is smaller than the actual number of taxonomic ranks.

Normalization of taxonomic ranks with the RED approach yielded values for 6 phyla, 14 classes, 41 orders, 90 families, and 247 genera (Fig. 5A; Supplementary Table 4). We found 85% of ranks fell within ± 0.1 of the median RED value for taxa at that rank, suggesting these ranks had comparable evolutionary divergence times. The only instance of a fungal rank that appears to be overclassified is the order Diaporthales, which contains plant pathogens (RED = 0.897; average RED value for other fungal orders = 0.752). All other instances that were outside the ± 0.1 RED interval concerned underclassification and appeared to be concentrated on specific lineages. Remarkably, nearly 40% (22 of 49, including 1 order, 5 families, 16 genera) of the underclassified ranks were within the Saccharomycotina subphylum of budding yeasts.

Other underclassified taxa included classes Chytridiomycetes (2 / 49), Tremellomycetes (2 / 49), and Agaricomycetes (4 / 49). The most underclassified lineage was order Zoopagales of Zoopagomycotina, whose RED value (= 0.309) was the lowest compared to other orders or classes included in our analysis. Since many Zoopagales are predacious or parasitic and non-culturable, all seven Zoopagales genomes have been sequenced using single cell sequencing methods^58^; thus, it is possible the low RED value in this lineage stems from the typically higher nucleotide base calling errors of single cell sequencing methods or from contamination. Moreover, it should be noted that the most serious instance of underclassification concerns the most well-sampled major lineage (Saccharomycotina). Thus, as the genomes of more species are sampled and added to the fungal phylogeny (especially from major lineages whose taxonomic diversity in not well represented in our phylogeny), it is possible that examination of RED values reveals further instances in the fungal tree of life where classification is not on par with evolutionary divergence.

Taken together, these results suggest that the current fungal taxonomy classification is largely concordant with our understanding of fungal phylogeny and evolutionary divergence. However, our results also identify lineages, such as Saccharomycotina, where taxonomic rank assignment appears to not truly reflect the observed levels of evolutionary divergence (compared to assignments in the rest of the fungal kingdom). Thus, an examination of the relative evolutionary divergence of fungal taxonomic groups of equivalent rank showed that they are unevenly applied across the kingdom, reducing the utility of taxonomy for comparative biology. Interestingly, our results show that the RED values and relative divergence time estimates by existing relaxed-clock models are significantly correlated (Fig. 5B), suggesting that both methods are interchangeable and can be used to compare taxonomy ranks in a phylogeny-informed way.

### Conclusion

Fungi have undergone extensive diversification into numerous ecological roles, morphological forms, and genomic architectures over the last one billion years (Fig. 1). Resolving relationships among major groups of the fungal tree has proven challenging due to the lack of data from organisms spanning fungal diversity and the relative paucity of phylogenomic studies for the entire kingdom. By synthesizing data from more than fifteen hundred publicly available genomes, we provide a robust phylogenetic framework to explore fungal evolution and examine sources of conflict and support for the backbone of the fungal phylogeny.

We find that most parts of the fungal phylogeny are robustly resolved with our 290 gene data set, but a handful of challenging branches remain unresolved. We provide evidence that some of these relationships may actually reflect genuine instances of ancient evolutionary diversification events, or hard polytomies, such as those among subphyla in Basidiomycota. In contrast, other unresolved relationships likely stem from the relatively poor taxon and / or gene sampling of several fungal phyla, suggesting that improving the resolution of the fungal phylogeny will require continued efforts to sample genomes spanning the diversity of the fungal kingdom. This inference is further supported by the results of our examination of concatenation- and coalescent-based phylogenies from several different data matrices that vary in their gene and taxon occupancy, which also suggests that the elucidation of these unresolved relationships will likely require substantial additional data and analyses. In the case of the monophyly of the Zygomycota, we show that the distinction between a phylogenomic analysis recovering monophyly versus paraphyly rests on a handful of genes. As fungal phylogenomic analyses improve their gene and taxon sampling, it is important to be aware that while the latest genome-scale phylogenies represent the currently best supported hypotheses, they are always potentially subject to revision and improvement.

Finally, our study presents a novel examination of the relationship between the current state of taxonomic classification in fungi and genomic evolutionary divergence. While fungal taxonomy broadly reflects evolutionary divergence, we identified instances of specific lineages, such as the subphylum Saccharomycotina, where the lack of correspondence hinders the utility of taxonomy as a yardstick for comparative biology. In conclusion, the generation and analyses of a phylogenomic data matrix from 1,644 species spanning the diversity of the kingdom establishes an integrated and robust phylogenetic framework for studying the evolution of the fungal kingdom.

## Methods

### Taxon sampling

All 1,679 fungal genomes were downloaded from NCBI and only one representative genome from every species was included (last accession date: January 30, 2020). Moreover, the genomes of 28 outgroup taxa (11 representative taxa from Holozoa and 17 representative taxa from Metazoa) were downloaded from Ensembl or NCBI (Last accession date: January 1, 2020). The outgroups were selected based on the current understanding of Opisthokonta phylogeny^29,30^. NCBI taxonomy, strain ID, and source information in this study are also provided in Supplementary Table 1.

### Quality assessment

To assess the qualities of the genome assemblies of the 1,679 fungal genomes we used the Benchmarking Universal Single-Copy Orthologs (BUSCO), version 2.02.1^31^ and the Fungi odb9 database (Last accession date: January 15, 2020). Briefly, BUSCO uses a consensus sequence built from a hidden Markov model-based alignment of orthologous sequences derived from 85 different fungal species using HMMER, version 3.1b2^59^, as a query in tBLASTn^60^ to search an individual genome. A total of 290 predefined orthologs (referred to as fungal BUSCO genes) were used. To examine the presence of each BUSCO gene in a genome, gene structure was predicted using AUGUSTUS, version 2.5.5^61^, with default parameters, from the nucleotide coordinates of putative genes identified using BLAST and then aligned to the HMM alignment of the same BUSCO gene. Genes were considered “single-copy” if there was only one complete predicted gene present in the genome, “duplicated” if there were two or more complete predicted genes for one BUSCO gene, “fragmented” if the predicted gene was shorter than 95% of the aligned sequence lengths from the 85 different fungal species, and “missing” if there was no predicted gene. For each genome, the fraction of single-copy BUSCO genes present corresponded to the completeness of each genome. To minimize missing data and remove potential low-quality genomes, were retained only those genomes that contained 100 or more single-copy BUSCO genes. The final data set contained 1,644 fungi and 28 outgroup taxa (Supplementary Table 1).

### Phylogenomic data matrix construction

In addition to their use as a measure of genome completeness, BUSCO genes have also been widely used as markers for phylogenomic inference in diverse lineages^31^, especially in exploring fungi relationships^21–23,62^. Therefore, we used the BUSCO genes to generate the full data matrix (1,672 taxa / 290 genes), as well as 11 additional data matrices by subsampling subsets of taxa or BUSCO genes. We used these 12 data matrices to assess the stability of phylogenetic relationships and identify putative sources of error in our analyses (Supplementary Fig. 2).

#### (1) Full data matrix

To construct the full data matrix, we only included single-copy BUSCO genes for each species. For each BUSCO gene, we extracted individual nucleotide sequences that have the BUSCO gene present and translated to amino acid sequences with their corresponding codon usage for each taxon (CUG-Ser1, CUG-Ser2 clades in yeasts: NCBI genetic code 12; CUG-Ala clades in yeasts: NCBI genetic code 26; all others: NCBI standard genetic code 1). Each gene was aligned with MAFFT version 7.299^63^ with options “—auto – maxiterate 1000”. Ambiguously aligned regions were removed using trimAl version 1.4^64^ with the “gappyout” option. The AA alignments of these 290 BUSCO genes, each of which has more than 50% of taxon occupancy, were then concatenated into the full data matrix, which contains 124,700 amino acid sites.

#### (2) Subset_Dikarya_taxa

Our taxon sampling is biased toward Ascomycota and Basidiomycota (Dikarya), especially in Saccharomycotina (332 taxa; 20.1 % total), Pezizomycotina (758 taxa; 46% total), and Agaricomycotina (321 taxa; 19.5% total). To discern the potential effects of biased taxon sampling (i.e., effects associated with the tree search algorithm spending most time in those parts of the tree that contain the largest numbers of taxa than in the other, less well sampled, parts of the tree), we subsampled one representative of each genus in Saccharomycotina (reducing their sampling from 332 taxa to 79), and one representative of each family in Pezizomycotina (758 -> 108 taxa) and in Agaricomycotina (321 -> 92 taxa). This sampling resulted in a data matrix with 540 taxa and 124,700 amino acid sites.

#### (3) Top_100_DVMC data matrix

This data matrix was constructed by retaining the top 100 BUSCO genes whose evolutionary rates were most “clock-like” (inferred by examining the degree of violation of a molecular clock^62^) and contains 51,494 amino acid sites (from all 1,672 taxa).

#### (4) Top_100_length data matrix

This data matrix was constructed by retaining the top 100 BUSCO genes with the longest alignment lengths after trimming and contains 75,529 amino acid sites (from all 1,672 taxa).

#### (5) Top100_low_LB data matrix

Long-Branch (LB) scores are widely used as a measurement for identifying genes that might be subject to long branch attraction^65^. LB scores were calculated for each species’ BUSCO gene using a customized python script (available at https://github.com/JLSteenwyk/Phylogenetic_scripts/blob/master/LB_score.py). This data matrix was constructed by retaining the top 100 BUSCO genes with the lowest average LB scores and contains 39,347 amino acid sites (from all 1,672 taxa).

#### (6) Top100_low_RCFV data matrix

This data matrix was constructed by retaining the 100 BUSCO genes with the lowest relative composition frequency variability (RCFV)^33^. The RCFV value for each gene was calculated following the protocols outlined by a previous study^21^. This data matrix contains 60,647 amino acid sites (from all 1,672 taxa).

#### (7) Top100_low_saturation data matrix

This data matrix was constructed by retaining the 100 BUSCO genes with the highest values of the slope of patristic distance – i.e., sum of the lengths of the branches that link two nodes in a tree – versus uncorrected p-distance (larger slope values denote lower levels of saturation than smaller values), which are thought to improve phylogenetic inference^66,67^. Slope values were measured by TreSpEx^33^. This data matrix contains 32,947 amino acid sites (from all 1,672 taxa).

#### (8) Top100_slow-evolving data matrix

This data matrix was constructed by retaining the 100 BUSCO genes with the lowest values of average pairwise patristic distance, which has previously been used to evaluate if fast-evolving genes bias phylogenetic inference^34,68^. The average patristic distance of each gene was measured by TreSpEx^33^. This data matrix contains 33,111 amino acid sites (from all 1,672 taxa).

#### (9) Top100_completeness data matrix

This data matrix was constructed by retaining the 100 BUSCO genes with the highest taxon occupancy. This data matrix contains 42,731 amino acid sites (from all 1,672 taxa).

#### (10) Top100_high_ABS data matrix

This data matrix was constructed by retaining the top 100 genes with the highest average bootstrap support (ABS) value of all internal branches on the gene tree in R package ape^69^, which has previously been shown to improve inference^70^. This data matrix contains 71,225 amino acid sites (from all 1,672 taxa).

#### (11) LB_taxa_removal

Long-Branch (LB) scores can also be used to identify taxa that might be subject to long branch attraction^65^. This data matrix was constructed by removal of 23 taxa with high LB score measured by a customized python script (https://github.com/JLSteenwyk/Phylogenetic_scripts/blob/master/LB_score.py). All 23 removed taxa were from the Microsporidia lineage. This removal resulted in a data matrix with 1,649 taxa and 124,700 amino acid sites.

#### (12) Rogue_taxa_removal

This data matrix was constructed by pruning 33 taxa that varied in their placement between analyses of the full data matrix by concatenation-based single model and coalescence using RogueNaRok^71^. This removal resulted in a data matrix with 1,639 taxa and 124,700 amino acid sites.

### Phylogenomic analyses

For the full data matrix as well as for each of these 11 data matrices constructed above, we used three different approaches to infer the fungal phylogeny: (1) the concatenation (i.e. supermatrix) approach with a single model or partition, (2) the concatenation approach with data-partitioning by gene, and (3) the multi-species coalescent-based approach that used the individual gene trees to construct the species phylogeny. All phylogenetic analyses were performed using IQ-TREE, version 1.6.8^72^, which has previously been shown to consistently perform well in analyses of phylogenomic data in a maximum likelihood (ML) framework^73^.

### Concatenation-based approach without and with data-partitioning

For concatenation-based analyses using a single model, we used the LG+G4 model^74^ because it was the best-fitting model for 89% of 290 gene trees. For analyses with data-partitioning by gene we used the best-fitting model for each gene (see coalescent-based approach section). Two independent runs were employed in all data matrices and the topological robustness of each gene tree was evaluated by 1,000 ultrafast bootstrap replicates^42^. A single tree search for the full data matrix (290 genes / 1,672 taxa) with a single model required ∼ 4,620 CPU hours.

### Coalescent-based approach

Individual gene trees were inferred using IQ-TREE, version 1.6.8 with an automatic detection for the best-fitting model with “-MFP” option using ModelFinder^75^ under the Bayesian information criterion (BIC). For each gene tree, we conducted 5 independent tree searches to obtain the best-scoring ML tree with “-runs 5” option. The topological robustness of each gene tree was evaluated by 1000 ultrafast bootstrap replicates.

To account for gene tree heterogeneity by taking incomplete lineage sorting (ILS) into account, we used the individual ML gene trees to infer the coalescent-based species tree using ASTRAL-III version 5.1.1^76^ for each data matrix. We applied contraction filters (BS < 33) such that poorly supported bipartitions within each gene tree were collapsed to polytomies, an approach recently suggested to improve the accuracy of ASTRAL^47^. The topological robustness was evaluated using the local posterior probability (LPP).

### Quantification of incongruence

From the set of 12 data matrices (the full one and 11 subsampled ones) and 3 analyses (concatenation with single model, concatenation with data-partitioning, and coalescence), we expect a total of 36 phylogenies. Data matrices 2, 11, and 12 have different sets of taxa that have been removed, so they cannot be straightforwardly compared to the rest of the data matrices, which contain the full set of taxa. To reduce the burden of computation (each tree search required thousands of CPU hours), we did not perform concatenation-based data-partitioning analyses for data matrices 1, 11 and 12. Thus, a total of 33 phylogenetic trees were compared.

For the 33 species phylogenies inferred from the 12 data matrices (12 from concatenation-based single model analyses, 9 from concatenation-based data-partitioning analyses, and 12 from coalescent-based analyses), we quantified the degree of incongruence for every internode by considering all prevalent conflicting bipartitions among individual ML gene trees^70,77^ using the “compare” function in Gotree version 1.13.6 (https://github.com/evolbioinfo/gotree).

### Polytomy test

To examine the support in individual gene trees for contentious bipartitions (and the alternative, conflicting bipartitions) and potentially identify evidence for hard polytomies of major fungal lineages, we used the polytomy test in ASTRAL, version 1.6.8^47^. The test evaluates whether a polytomy can be rejected by examining the frequencies of the three alternative quartet tree topologies in a set of trees. In our case, we used all gene trees as input for the calculation of the frequencies of the three alternative quartet trees for bipartitions of interest. In all cases, we used a P value cutoff of < 0.05 to reject the null hypothesis of a polytomy (see Fig. 4 for eight tested hypotheses). We used scripts available at https://github.com/smirarab/1kp/tree/master/scripts/hypo-test. We used pos-for-hyp-4-11-2.sh (-t 4 option) and quart-for-hyp-4-11-2.sh (-t 8 option) to compute the posterior probabilities for all three alternative topologies of a given quartet. To evaluate the discordance of gene trees in our single-copy gene data set, we used the *Q* value in ASTRAL to display the percentages of quartets in gene trees in support of the topology inferred by concatenation (q1) as well as the other two possible alternative topologies (q2 and q3); We used poly-for-hyp-4-11-02.sh to compute the p-value for a hard polytomy under the null hypothesis using ASTRAL (-t 10 option).

### Quantification of the distribution of phylogenetic signal

To investigate the distribution of phylogenetic signal of whether Zygomycota are monophyletic or paraphyletic, we considered two data matrices that had different topologies between ML analyses. To save computation time, we used the subset Dikarya data matrix (#2) since it has essentially the same topology as the full data matrix but has many fewer taxa. We also analyzed the Spatafora2016_46taxa_192genes data matrix from a previous study that recovered the paraphyly of Zygomycota^18^. We examined two hypotheses: Zygomycota-monophyly (T1) and Zygomycota-paraphyly (T2: Zoopagomycota sister to Dikarya + Mucoromycota). For ML analysis in each data matrix, site-wise likelihood scores were inferred for both hypotheses using IQ-TREE, version 1.6.8 (option -g) with the LG+G4 model. The two different phylogenetic trees passed to IQ-TREE (via -z) were the tree where Zygomycota is monophyletic and a tree modified to have Zoopagomycota placed as the sister to Dikarya + Mucoromycota. The numbers of genes and sites supporting each hypothesis were calculated from IQ-TREE output and Perl scripts from a previous study^24^. By calculating gene-wise log-likelihood scores between T1 and T2 for every gene, we considered a gene with an absolute value of log-likelihood difference of two as a gene with strong (|ΔlnL| > 2) or weak (|ΔlnL| < 2) phylogenetic signal as done in a previous study^78^.

### RED index

To evaluate whether fungal taxonomy is consistent with evolutionary genomic divergence, we calculated relative evolutionary divergence (RED) values from the annotated tree inferred from the full data matrix using concatenation with a single model by PhyloRank (v0.0.37; https://github.com/dparks1134/PhyloRank/), as described previously^57^. Briefly, the NCBI taxonomy associated with every fungal genome was obtained from the NCBI Taxonomy FTP site on January 17, 2020. PhyloRank linearly interpolates the RED values of every internal node according to lineage-specific rates of evolution under the constraints of the root being defined as zero and the RED of all present taxa being defined as one^57,79^. The RED intervals for each rank were defined as the median RED value ± 0.1 to serve as a guide for the normalization of taxonomic ranks from genus to phylum.

We also compared RED values to relative time divergence under a relaxed-molecular clock model for every taxonomic rank from genus to phylum, since both methods are based on inferring lineage-specific rates of evolution. We used the RelTime algorithm employed in the command line version of MEGA7^80^ since it is computationally much less demanding than Bayesian tree-dating methods. We conducted divergence time estimation using the full data matrix with the same ML tree that we used for the RED analysis (see above) without fossil calibrations. Correlation between the RED values and relative divergence time estimated by RelTime was calculated using Pearson’s correlation coefficient using the cor.test function in R package stats v.3.6.2^81^.

## Data and code availability

Upon publication, all genome assemblies will become publicly available in the Zenodo repository. Similarly, all scripts, data matrices, and phylogenetic trees will become publicly available at Figshare.

## Acknowledgments

We thank members of the Rokas laboratory for discussions and comments, Phil Hugenholtz for encouraging us to use relative evolutionary distance to examine the correspondence between genome sequence divergence and taxonomic rank across fungi, and Donovan Parks for initial help with getting relative evolutionary distance analyses working. This work was conducted in part using the resources of the Advanced Computing Center for Research and Education (ACCRE) at Vanderbilt University and Yale Center for Research Computing (Farnam HPC cluster) for use of the research computing infrastructure. Yuanning Li was partially supported by a scholarship from the China Scholarship Council (CSC) for studying and living abroad. This work was supported by the National Science Foundation (DEB-1442113 to A.R.; DEB-1442148 to C.T.H.; DEB-1929738 and DEB-1441677 to T.Y.J.; DEB-1557110 and DEB-1441715 to J.E.S; DEB-1441604 to J.W.S), in part by the DOE Great Lakes Bioenergy Research Center (DOE Office of Science BER DE-FC02-07ER64494), and the USDA National Institute of Food and Agriculture (Hatch project 1003258 to C.T.H.; Hatch project CA-R-PPA-5062-H to J.E.S.). C.T.H. is a Pew Scholar in the Biomedical Sciences and a H. I. Romnes Faculty Fellow, supported by the Pew Charitable Trusts and Office of the Vice Chancellor for Research and Graduate Education with funding from the Wisconsin Alumni Research Foundation, respectively. X.X.S. was supported by the start-up grant from the “Hundred Talents Program” at Zhejiang University and the Fundamental Research Funds for the Central Universities (2020QNA6019). J.E.S. is a Fellow in the CIFAR program Fungal Kingdom: Threats and Opportunities. J.L.S. and A.R. were supported by the Howard Hughes Medical Institute through the James H. Gilliam Fellowships for Advanced Study program. A.R. received additional support from the Guggenheim Foundation and the Burroughs Wellcome Fund.

## Supplementary Figure Legends

**Supplementary Figure 1. Current consensus of evolutionary relationships of major lineages within kingdom Fungi**. Phyla not sampled in this study are shown in red font.

**Supplementary Figure 2. Relationships between the 12 data matrices analyzed in this study**. Data matrices with taxon-based filtering are in purple boxes and those with gene-based filtering are in green boxes. The number for each data matrix corresponds to its number in the Methods section. See Methods for further information on each data matrix and filtering strategy used to generate it.

**Supplementary Figure 3. The genome-scale phylogeny of 1**,**644 species in the fungal kingdom**. The tree of the 1,644 fungal species and 28 outgroups was reconstructed from the maximum likelihood concatenation analysis of 290 single-copy BUSCO genes under a single LG+G4 model (*lnL* = -78287339.984). All internal branches were supported with 100% bootstrap value unless otherwise noted. See also Figure 3A and Supplementary Table 1.

**Supplementary Figure 4. Heatmap of topological similarities for all pairwise comparisons among the phylogenies reconstructed from analyses of 12 different data matrices using three different approaches (concatenation under a single partition, concatenation under gene-based partitioning, and coalescence)**. The topological congruence between each pair of phylogenies was calculated using Gotree. The size and color of the squares represents the degree of congruence as measured by percentage. Results from data matrices 2, 11, and 12 are not shown here since they have different sets of taxa that have been removed.

**Supplementary Figure 5. Phylogeny of 1**,**639 fungal species from the Rogue_taxa_removal data matrix**. The topology shown was obtained from maximum likelihood analysis of a concatenated data matrix of 290 genes under a single LG+G4 model (*lnL* = -76877622.807). All internal branches were supported with 100% bootstrap values unless otherwise noted. Each tip corresponds to the class-level ranking derived from NCBI taxonomy (except for subphylum Saccharomycotina, where each tip corresponds to each one of the 12 major clades to reflect the current understanding of Saccharomycotina phylogeny^21^).

**Supplementary Figure 6. Phylogeny of 1**,**644 fungal species from the Top100_slow_evolving data matrix under a single LG+G4 model**. The topology shown was obtained from maximum likelihood analysis of a concatenated data matrix of 290 genes under a single LG+G4 model (*lnL* = -13426586.414). All internal branches were supported with 100% bootstrap values unless otherwise noted. Each tip corresponds to the class-level ranking derived from NCBI taxonomy (except for subphylum Saccharomycotina, where each tip corresponds to each one of the 12 major clades to reflect the current understanding of Saccharomycotina phylogeny^21^).

**Supplementary Figure 7. Distribution of phylogenetic signal for two alternative hypotheses on the Zygomycota lineage**. The two alternative hypotheses are: Mucoromycota is sister to Zoopagomycota (Zygomycota-monophyly; T1 Orange), Mucoromycota is sister to Dikarya (Zygomycota-paraphyly; T2 Green). Proportions of genes supporting each of two alternative hypotheses in the Spatafora2016_46taxa_192genes and Subset_Dikaya data matrices. The GLS values for each gene in each data matrix are provided in Supplementary Table 5. We considered a gene with an absolute value of log-likelihood difference of two as a gene with strong (|ΔlnL| > 2) or weak (|ΔlnL| < 2) phylogenetic signal.

## Supplementary Tables

**Supplementary Table 1**. Taxonomy, strain ID, and source information of 1,644 fungi and 28 outgroup genomes.

**Supplementary Table 2**. Gene statistics, taxon occupancies, and annotations used for sensitive analyses of 290 fungi BUSCO genes.

**Supplementary Table 3**. An examination of monophyletic lineages in different taxonomic levels of class, subphylum, and phylum.

**Supplementary Table 4**. Detailed results of RED values for each taxa based on NCBI taxonomy information.

**Supplementary Table 5**. Distribution of phylogenetic signal for two alternative hypotheses on the Zygomycota lineage.

## References

1. Blackwell, M. The Fungi: 1, 2, 3… 5.1 million species? Am. J. Bot. 98, 426–438 (2011).

2. Hawksworth, D. L. & Lücking, R. Fungal Diversity Revisited: 2.2 to 3.8 Million Species. Microbiol Spectr 5, (2017).

3. Heitman, J. et al. The Fungal Kingdom. (John Wiley & Sons, 2017).

4. Lutzoni, F. et al. Contemporaneous radiations of fungi and plants linked to symbiosis. Nat. Commun. 9, 5451 (2018).

5. Naranjo-Ortiz, M. A. & Gabaldón, T. Fungal evolution: major ecological adaptations and evolutionary transitions. Biol. Rev. Camb. Philos. Soc. 94, 1443–1476 (2019).

6. Timothy Y. James, Jason E. Stajich, Chris Todd Hittinger, Antonis Rokas. Toward a Fully Resolved Fungal Tree of Life. Annu. Rev. Microbiol. 74, (2020).

7. Wijayawardene, N. N. et al. Outline of Fungi and fungus-like taxa. Embrapa Milho e Sorgo-Artigo em periódico indexado (ALICE) (2020).

8. Spatafora, J. W. et al. The Fungal Tree of Life: from Molecular Systematics to Genome-Scale Phylogenies. Microbiol Spectr 5, (2017).

9. Torruella, G. et al. Global transcriptome analysis of the aphelid supports the phagotrophic origin of fungi. Commun Biol 1, 231 (2018).

10. Bass, D. et al. Clarifying the Relationships between Microsporidia and Cryptomycota. J. Eukaryot. Microbiol. 65, 773–782 (2018).

11. Ebersberger, I. et al. A consistent phylogenetic backbone for the fungi. Mol. Biol. Evol. 29, 1319–1334 (2012).

12. Chang, Y. et al. Phylogenomic Analyses Indicate that Early Fungi Evolved Digesting Cell Walls of Algal Ancestors of Land Plants. Genome Biol. Evol. 7, 1590–1601 (2015).

13. Torruella, G. et al. Phylogenomics Reveals Convergent Evolution of Lifestyles in Close Relatives of Animals and Fungi. Curr. Biol. 25, 2404–2410 (2015).

14. Hibbett, D. S. et al. A higher-level phylogenetic classification of the Fungi. Mycol. Res. 111, 509–547 (2007).

15. Schüβler, A., Schwarzott, D. & Walker, C. A new fungal phylum, the Glomeromycota: phylogeny and evolution* *Dedicated to Manfred Kluge (Technische Universität Darmstadt) on the occasion of his retirement. Mycol. Res. 105, 1413–1421 (2001).

16. James, T. Y. et al. Reconstructing the early evolution of Fungi using a six-gene phylogeny. Nature 443, 818–822 (2006).

17. Capella-Gutiérrez, S., Marcet-Houben, M. & Gabaldón, T. Phylogenomics supports microsporidia as the earliest diverging clade of sequenced fungi. BMC Biol. 10, 47 (2012).

18. Spatafora, J. W. et al. A phylum-level phylogenetic classification of zygomycete fungi based on genome-scale data. Mycologia 108, 1028–1046 (2016).

19. Varga, T. et al. Megaphylogeny resolves global patterns of mushroom evolution. Nature Ecology & Evolution vol. 3 668–678 (2019).

20. Kiss, E. et al. Comparative genomics reveals the origin of fungal hyphae and multicellularity. Nat. Commun. 10, 4080 (2019).

21. Shen, X.-X. et al. Tempo and Mode of Genome Evolution in the Budding Yeast Subphylum. Cell 175, 1533–1545.e20 (2018).

22. Shen, X. X., Steenwyk, J. L., LaBella, A. L. & Opulente, D. A. Genome-scale phylogeny and contrasting modes of genome evolution in the fungal phylum Ascomycota. bioRxiv (2020).

23. Shen, X.-X. et al. Reconstructing the Backbone of the Saccharomycotina Yeast Phylogeny Using Genome-Scale Data. G3 6, 3927–3939 (2016).

24. Shen, X.-X., Hittinger, C. T. & Rokas, A. Contentious relationships in phylogenomic studies can be driven by a handful of genes. Nat Ecol Evol 1, 126 (2017).

25. Grigoriev, I. V. et al. MycoCosm portal: gearing up for 1000 fungal genomes. Nucleic Acids Res. 42, (2014).

26. Hittinger, C. T. et al. Genomics and the making of yeast biodiversity. Curr. Opin. Genet. Dev. 35, (2015).

27. Haridas, S. et al. 101 Dothideomycetes genomes: A test case for predicting lifestyles and emergence of pathogens. Stud. Mycol. 96, 141–153 (2020).

28. Vesth, T. C. et al. Investigation of inter- and intraspecies variation through genome sequencing of Aspergillus section Nigri. Nat. Genet. 50, 1688–1695 (2018).

29. Torruella, G. et al. Phylogenetic relationships within the Opisthokonta based on phylogenomic analyses of conserved single-copy protein domains. Mol. Biol. Evol. 29, 531–544 (2012).

30. Brown, M. W., Spiegel, F. W. & Silberman, J. D. Phylogeny of the ‘Forgotten’ Cellular Slime Mold, Fonticula alba, Reveals a Key Evolutionary Branch within Opisthokonta. Mol. Biol. Evol. 26, 2699–2709 (2009).

31. Waterhouse, R. M. et al. BUSCO Applications from Quality Assessments to Gene Prediction and Phylogenomics. Mol. Biol. Evol. 35, 543–548 (2018).

32. Zdobnov, E. M. et al. OrthoDB v9.1: cataloging evolutionary and functional annotations for animal, fungal, plant, archaeal, bacterial and viral orthologs. Nucleic Acids Res. 45, D744–D749 (2017).

33. Struck, T. H. TreSpEx–-Detection of Misleading Signal in Phylogenetic Reconstructions Based on Tree Information. Evol. Bioinform. Online 10, EBO.S14239 (2014).

34. Kocot, K. M. et al. Phylogenomics of Lophotrochozoa with Consideration of Systematic Error. Syst. Biol. 66, 256–282 (2017).

35. Quandt, C. A. et al. The genome of an intranuclear parasite, Paramicrosporidium saccamoebae, reveals alternative adaptations to obligate intracellular parasitism. Elife 6, (2017).

36. Zajc, J. et al. Genome and transcriptome sequencing of the halophilic fungus Wallemia ichthyophaga: haloadaptations present and absent. BMC Genomics 14, 617 (2013).

37. Padamsee, M. et al. The genome of the xerotolerant mold Wallemia sebi reveals adaptations to osmotic stress and suggests cryptic sexual reproduction. Fungal Genet. Biol. 49, 217–226 (2012).

38. Zhao, R.-L. et al. A six-gene phylogenetic overview of Basidiomycota and allied phyla with estimated divergence times of higher taxa and a phyloproteomics perspective. Fungal Divers. 84, 43–74 (2017).

39. Choi, J. & Kim, S.-H. A genome Tree of Life for the Fungi kingdom. Proc. Natl. Acad. Sci. U. S. A. 114, 9391–9396 (2017).

40. Li, Y. et al. Feature Frequency Profile-based phylogenies are inaccurate. bioRxiv (2020) doi:10.1101/2020.06.28.176479.

41. James, T. Y. et al. Shared signatures of parasitism and phylogenomics unite Cryptomycota and microsporidia. Curr. Biol. 23, 1548–1553 (2013).

42. Hoang, D. T., Chernomor, O., von Haeseler, A., Minh, B. Q. & Vinh, L. S. UFBoot2: Improving the Ultrafast Bootstrap Approximation. Mol. Biol. Evol. 35, 518–522 (2018).

43. Rokas, A., Williams, B. L., King, N. & Carroll, S. B.Genome-scale approaches to resolving incongruence in molecular phylogenies.Nature 425, 798–804 (2003).

44. Zhang, C., Rabiee, M., Sayyari, E. & Mirarab, S. ASTRAL-III: polynomial time species tree reconstruction from partially resolved gene trees. BMC Bioinformatics 19, 153 (2018).

45. Gatesy, J. & Springer, M. S. Phylogenetic analysis at deep timescales: unreliable gene trees, bypassed hidden support, and the coalescence/concatalescence conundrum. Mol. Phylogenet. Evol. 80, 231–266 (2014).

46. Edwards, S. V. et al. Implementing and testing the multispecies coalescent model: A valuable paradigm for phylogenomics. Molecular Phylogenetics and Evolution vol. 94 447–462 (2016).

47. Sayyari, E. & Mirarab, S. Testing for Polytomies in Phylogenetic Species Trees Using Quartet Frequencies. Genes 9, (2018).

48. Ruiz-Herrera, J. & Ortiz-Castellanos, L. Cell wall glucans of fungi. A review. The Cell Surface vol. 5 100022 (2019).

49. Dee, J. M., Mollicone, M., Longcore, J. E., Roberson, R. W. & Berbee, M. L. Cytology and molecular phylogenetics of Monoblepharidomycetes provide evidence for multiple independent origins of the hyphal habit in the Fungi. Mycologia 107, 710–728 (2015).

50. Liu, Y. et al. Phylogenomic analyses predict sistergroup relationship of nucleariids and fungi and paraphyly of zygomycetes with significant support. BMC Evol. Biol. 9, 272 (2009).

51. Spatafora, J. W. et al. A phylum-level phylogenetic classification of zygomycete fungi based on genome-scale data. Mycologia 108, 1028–1046 (2016).

52. Sekimoto, S., Rochon, D. ‘ann, Long J. E., Dee, J. M. & Berbee, M. L. A multigene phylogeny of Olpidium and its implications for early fungal evolution. BMC Evol. Biol. 11, 331 (2011).

53. Gryganskyi, A. P. et al. Phylogenetic lineages in Entomophthoromycota. Persoonia 30, 94–105 (2013).

54. Tabima, J. F. et al. Phylogenomic analyses of non-Dikarya fungi supports horizontal gene transfer driving diversification of secondary metabolism in the amphibian gastrointestinal symbiont, Basidiobolus. doi:10.1101/2020.04.08.030916.

55. He, M.-Q. et al. Notes, outline and divergence times of Basidiomycota. Fungal Divers. 99, 105–367 (2019).

56. Prasanna, A. N. et al. Model Choice, Missing Data, and Taxon Sampling Impact Phylogenomic Inference of Deep Basidiomycota Relationships. Syst. Biol. 69, 17–37 (2020).

57. Parks, D. H. et al. A standardized bacterial taxonomy based on genome phylogeny substantially revises the tree of life. Nat. Biotechnol. 36, 996–1004 (2018).

58. Davis, W. J. et al. Genome-scale phylogenetics reveals a monophyletic Zoopagales (Zoopagomycota, Fungi). Mol. Phylogenet. Evol. 133, 152–163 (2019).

59. Zhang, Z. & Wood, W. I. A profile hidden Markov model for signal peptides generated by HMMER. Bioinformatics 19, 307–308 (2003).

60. Gertz, E. M., Yu, Y.-K., Agarwala, R., Schäffer, A. A. & Altschul, S. F. Composition-based statistics and translated nucleotide searches: improving the TBLASTN module of BLAST. BMC Biol. 4, 41 (2006).

61. Stanke, M. et al. AUGUSTUS: ab initio prediction of alternative transcripts. Nucleic Acids Res. 34, W435–W439 (2006).

62. Steenwyk, J. L., Shen, X.-X., Lind, A. L., Goldman, G. H. & Rokas, A. A Robust Phylogenomic Time Tree for Biotechnologically and Medically Important Fungi in the Genera Aspergillus and Penicillium. MBio 10, (2019).

63. Katoh, K. & Standley, D. M. MAFFT multiple sequence alignment software version 7: improvements in performance and usability. Mol. Biol. Evol. 30, 772–780 (2013).

64. Capella-Gutiérrez, S., Silla-Martínez, J. M. & Gabaldón, T. trimAl: a tool for automated alignment trimming in large-scale phylogenetic analyses. Bioinformatics 25, 1972–1973 (2009).

65. Felsenstein, J. Cases in which Parsimony or Compatibility Methods will be Positively Misleading. Syst. Biol. 27, 401–410 (1978).

66. Struck, T. H., Nesnidal, M. P., Purschke, G. & Halanych, K. M. Detecting possibly saturated positions in 18S and 28S sequences and their influence on phylogenetic reconstruction of Annelida (Lophotrochozoa). Mol. Phylogenet. Evol. 48, 628–645 (2008).

67. Kocot, K. M. et al. Phylogenomics of Lophotrochozoa with Consideration of Systematic Error. Syst. Biol. 66, 256–282 (2017).

68. Whelan, N. V., Kocot, K. M., Moroz, L. L. & Halanych, K. M. Error, signal, and the placement of Ctenophora sister to all other animals. Proc. Natl. Acad. Sci. U. S. A. 112, 5773–5778 (2015).

69. Paradis, E. & Schliep, K. ape 5.0: an environment for modern phylogenetics and evolutionary analyses in R. Bioinformatics 35, 526–528 (2019).

70. Salichos, L. & Rokas, A. Inferring ancient divergences requires genes with strong phylogenetic signals. Nature 497, 327–331 (2013).

71. Aberer, A. J., Krompass, D. & Stamatakis, A. Pruning rogue taxa improves phylogenetic accuracy: an efficient algorithm and webservice. Syst. Biol. 62, 162–166 (2013).

72. Minh, B. Q. et al. IQ-TREE 2: New models and efficient methods for phylogenetic inference in the genomic era. Mol. Biol. Evol. (2020) doi:10.1093/molbev/msaa015.

73. Zhou, X., Shen, X.-X., Hittinger, C. T. & Rokas, A. Evaluating Fast Maximum Likelihood-Based Phylogenetic Programs Using Empirical Phylogenomic Data Sets. Mol. Biol. Evol. 35, 486–503 (2018).

74. Le, S. Q., Lartillot, N. & Gascuel, O. Phylogenetic mixture models for proteins. Philos. Trans. R. Soc. Lond. B Biol. Sci. 363, 3965–3976 (2008).

75. Kalyaanamoorthy, S., Minh, B. Q., Wong, T. K. F., von Haeseler, A. & Jermiin, L. S. ModelFinder: fast model selection for accurate phylogenetic estimates. Nat. Methods 14, 587–589 (2017).

76. Mirarab, S. et al. ASTRAL: genome-scale coalescent-based species tree estimation. Bioinformatics 30, i541–8 (2014).

77. Salichos, L., Stamatakis, A. & Rokas, A. Novel information theory-based measures for quantifying incongruence among phylogenetic trees. Mol. Biol. Evol. 31, 1261–1271 (2014).

78. Smith, S. A., Walker-Hale, N., Walker, J. F. & Brown, J. W. Phylogenetic conflicts, combinability, and deep phylogenomics in plants. Syst. Biol. 69, 579–592 (2020).

79. Rinke, C. et al. A rank-normalized archaeal taxonomy based on genome phylogeny resolves widespread incomplete and uneven classifications. bioRxiv 2020.03.01.972265 (2020) doi:10.1101/2020.03.01.972265.

80. Kumar, S., Stecher, G. & Tamura, K. MEGA7: Molecular Evolutionary Genetics Analysis Version 7.0 for Bigger Datasets. Mol. Biol. Evol. 33, 1870–1874 (2016).

81. Team, R. C., Team, M. R. C., Suggests, M. & Matrix, S. Package stats. The R Stats Package (2018).

