## Supplementary Figures for "A genome-scale phylogeny of Fungi; insights into early evolution, radiations, and the relationship between taxonomy and phylogeny"

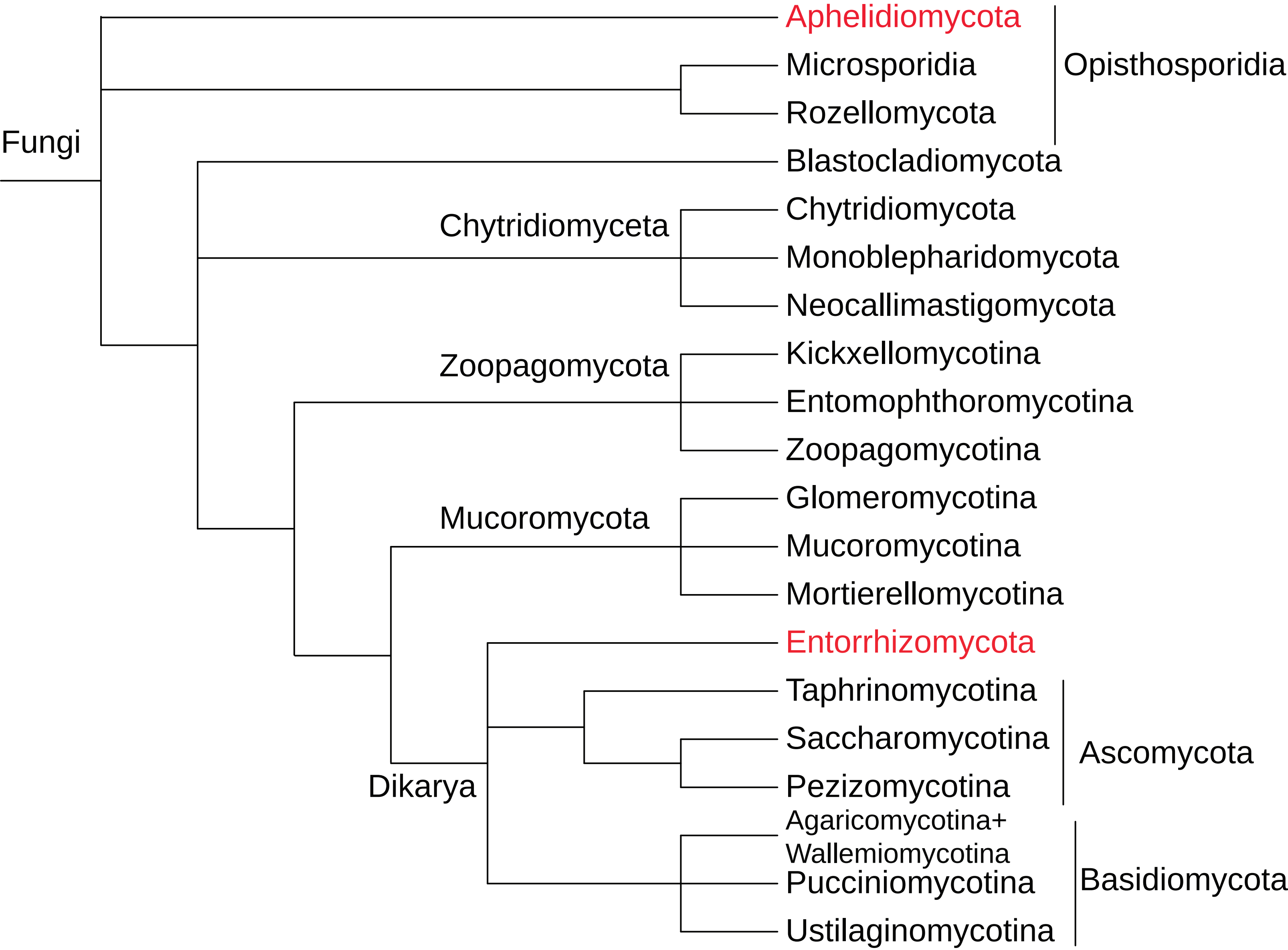


**Supplementary Figure 1. Current consensus of evolutionary relationships of major lineages within kingdom Fungi.** Phyla not sampled in this study are shown in red font.

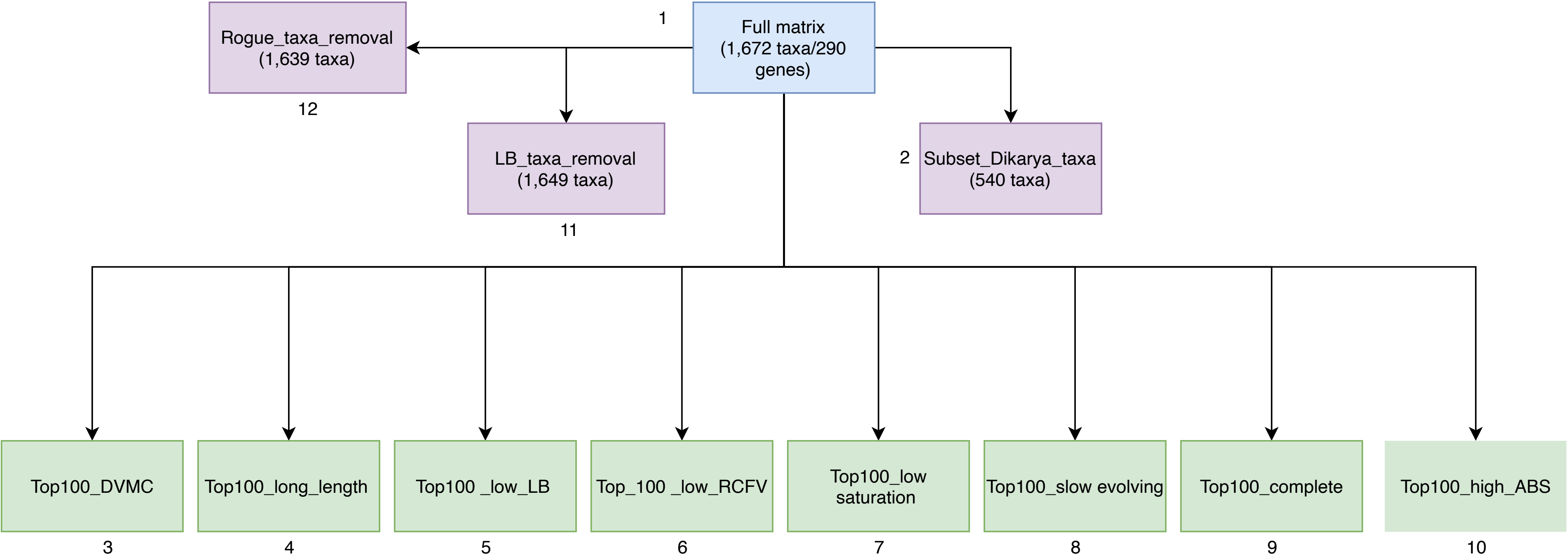


**Supplementary Fig. 2. Relationships between the 12 data matrices analyzed in this study.** Data matrices with taxon-based filtering are in purple boxes and those with gene-based filtering are in green boxes. The number for each data matrix corresponds to its number in the Methods section. See Methods for further information on each data matrix and filtering strategy used to generate it.

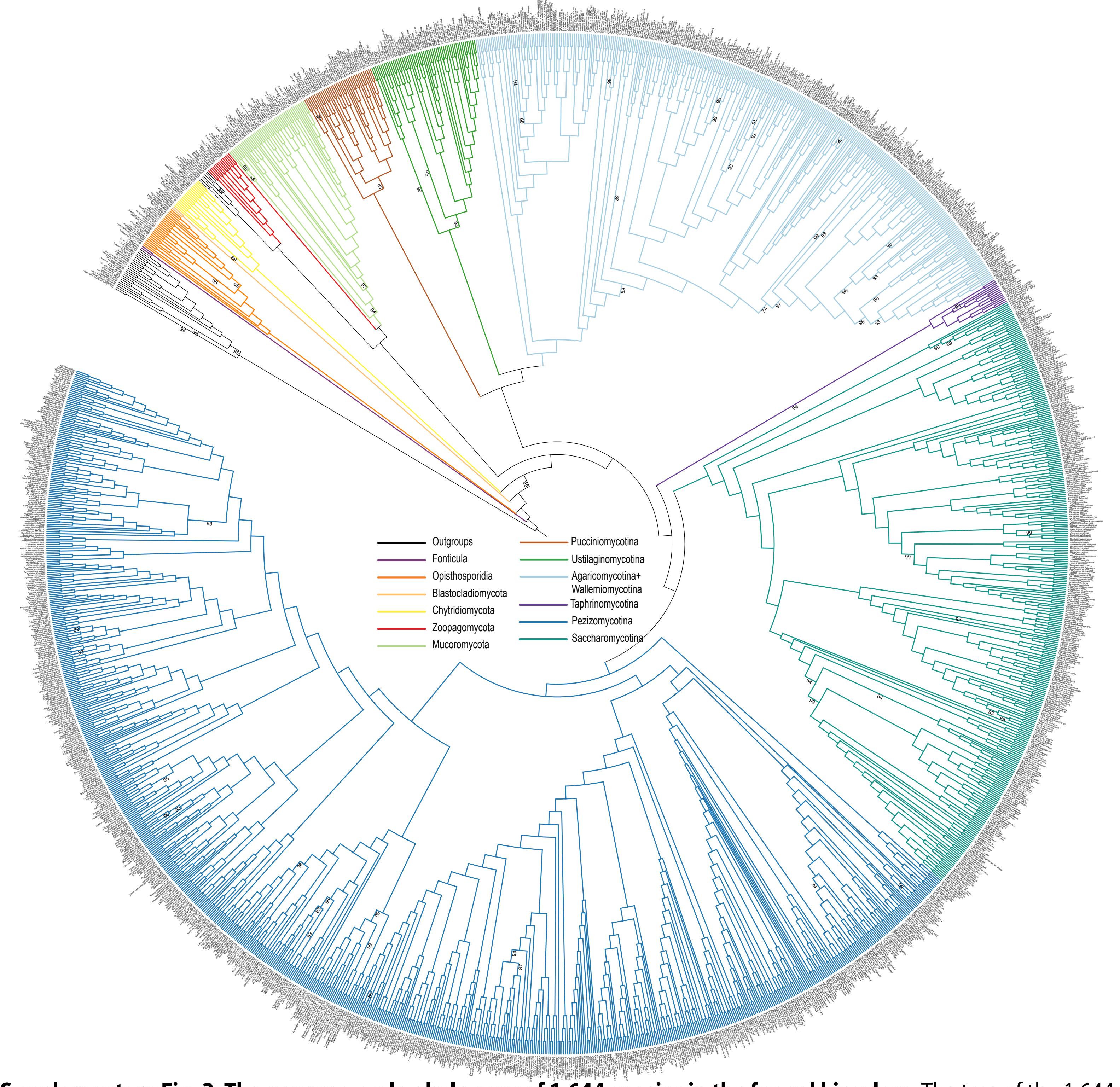


**Supplementary Fig. 3. The genome-scale phylogeny of 1,644 species in the fungal kingdom.** The tree of the 1,644 fungal species and 28 outgroups was reconstructed from the maximum likelihood concatenation analysis of 290 single-copy BUSCO genes under a single LG+G4 model (lnL = -78287339.984). All internal branches were supported with 100% bootstrap value unless otherwise noted. See also Figure 3A and Supplementary Table 1.

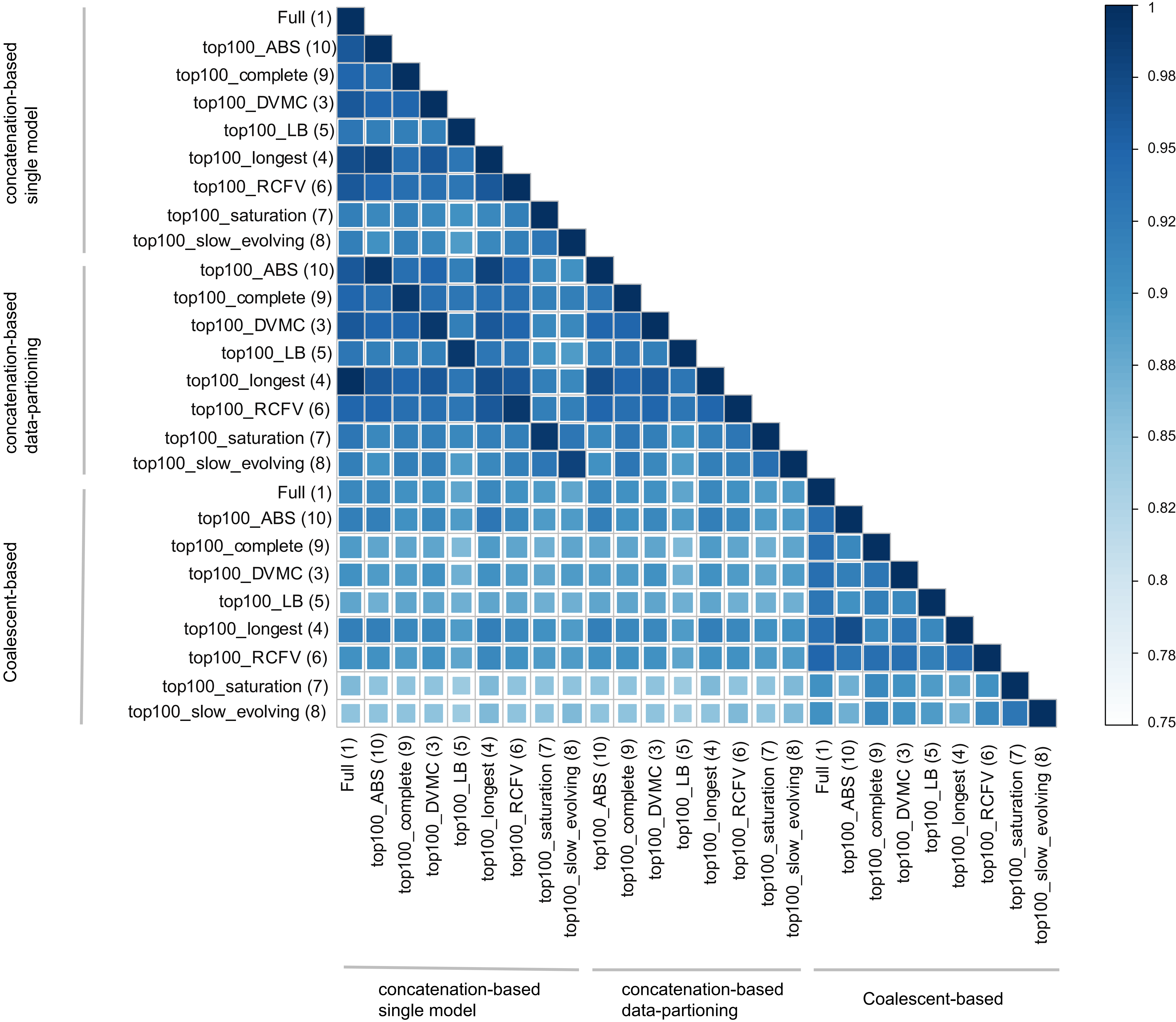


**Supplementary Fig. 4. Heatmap of topological similarities for all pairwise comparisons among the phylogenies reconstructed from analyses of 12 different data matrices using three different approaches (concatenation under a single partition, concatenation under gene-based partitioning, and coalescence).** The topological congruence between each pair of phylogenies was calculated using Gotree. The size and color of the squares represents the degree of congruence as measured by percentage. Results from data matrices 2, 11, and 12 are not shown here since they have different sets of taxa that have been removed.

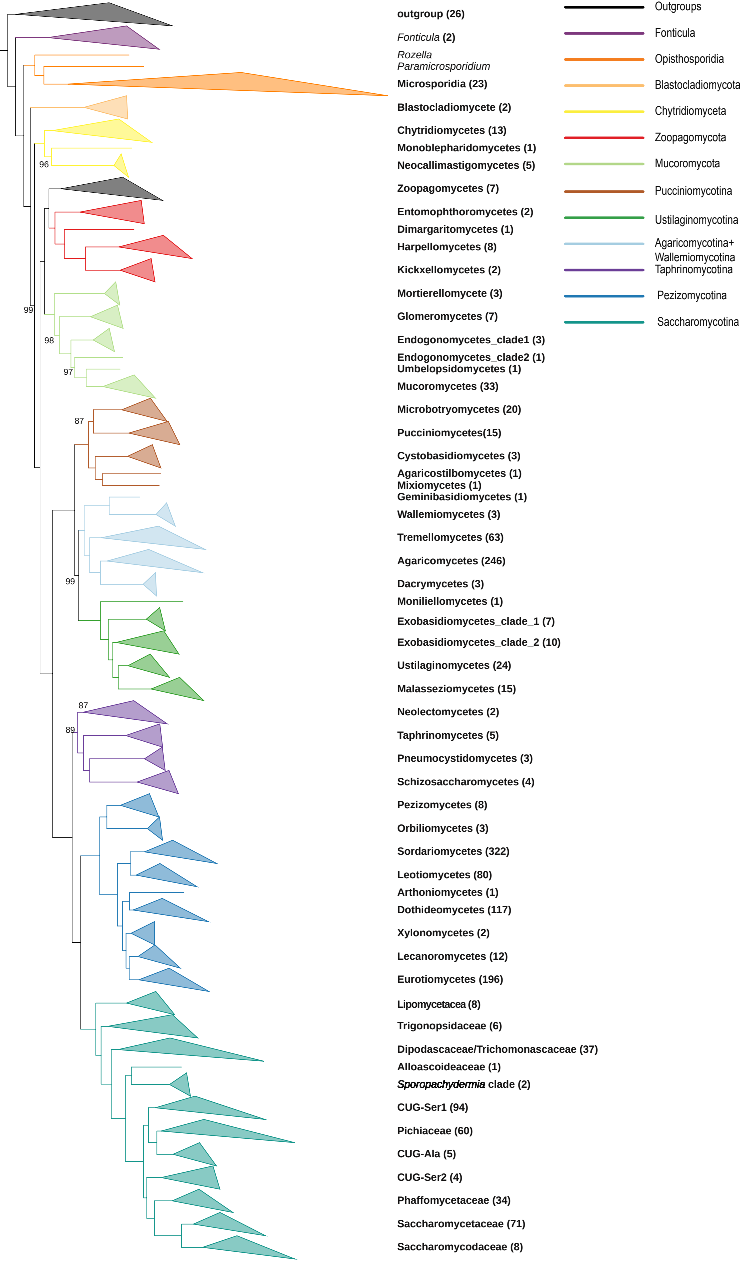


**Supplementary Fig. 5. Phylogeny of 1,639 fungal species from the Rogue_taxa_removal data matrix.** The topology shown was obtained from maximum likelihood analysis of a concatenated data matrix of 290 genes under a single LG+G4 model (lnL = -76877622.807). All internal branches were supported with 100% bootstrap values unless otherwise noted. Each tip corresponds to the class-level ranking derived from NCBI taxonomy (except for subphylum Saccharomycotina, where each tip corresponds to each one of the 12 major clades to reflect the current understanding of Saccharomycotina phylogeny[21]).

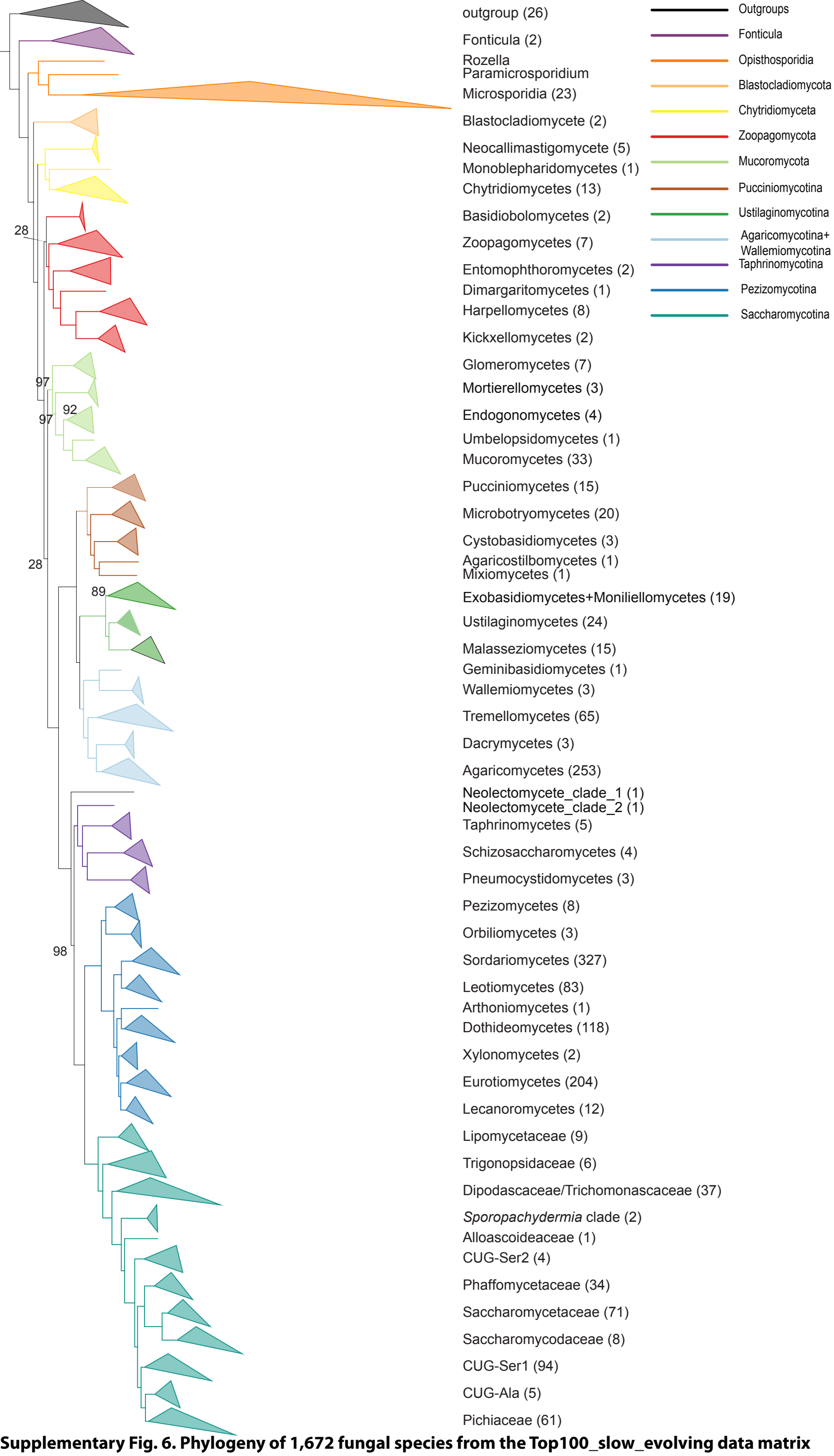


**Supplementary Fig. 6. Phylogeny of 1,672 fungal species from the Top100_slow_evolving data matrix under a single LG+G4 model.** The topology shown was obtained from maximum likelihood analysis of a concatenated data matrix of 290 genes under a single LG+G4 model (lnL = -13426586.414). All internal branches were supported with 100% bootstrap values unless otherwise noted. Each tip corresponds to the class-level ranking derived from NCBI taxonomy (except for subphylum Saccharomycotina, where each tip corresponds to each one of the 12 major clades to reflect the current understanding of Saccharomycotina phylogeny[21]).

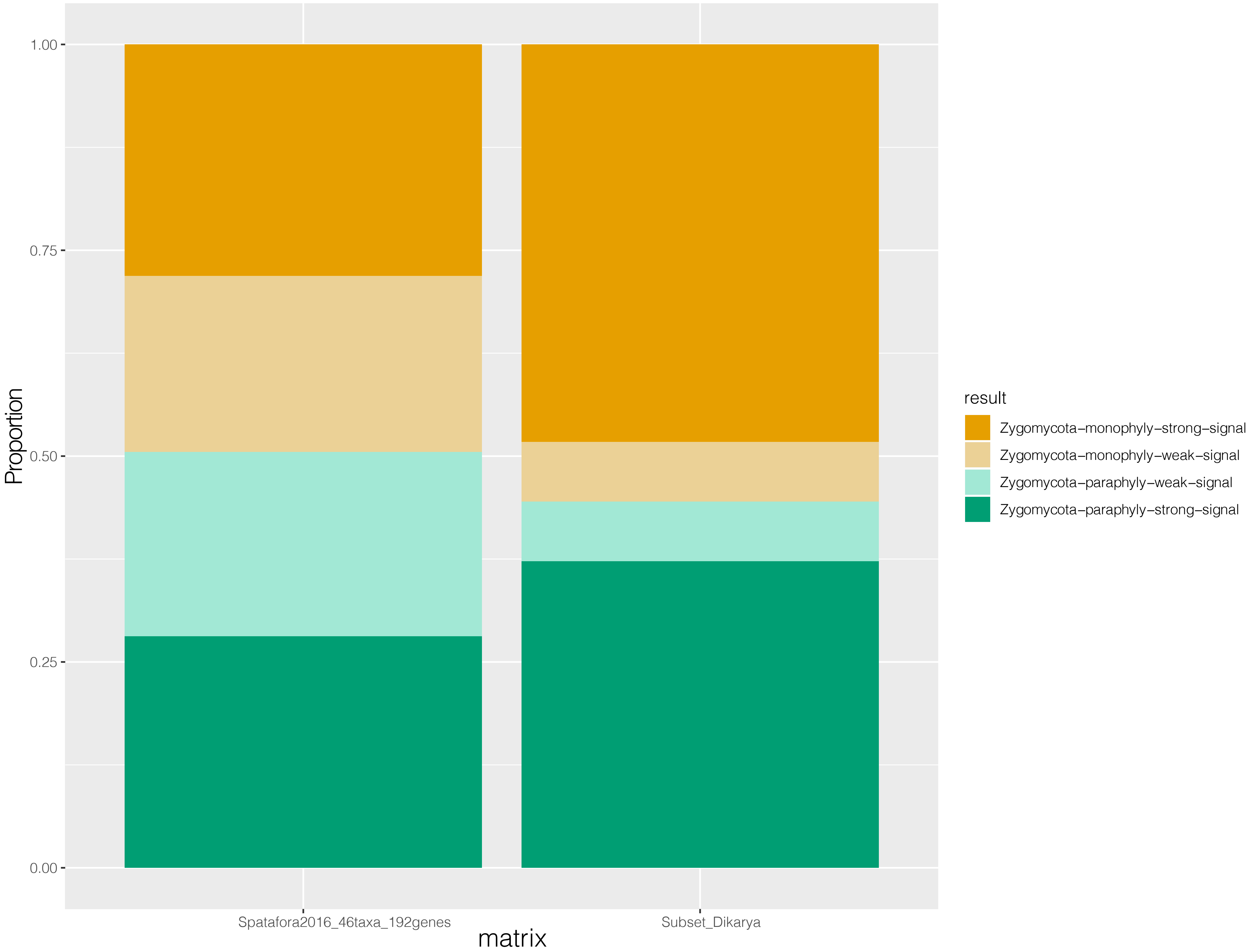


**Supplementary Fig. 7 . Distribution of phylogenetic signal for two alternative hypotheses on the Zygomycota lineage.** The two alternative hypotheses are: Mucoromycota is sister to Zoopagomycota (Zygomycota-monophyly; T1 Orange), Mucoromycota is sister to Dikarya (Zygomycota-paraphyly; T2 Green). Proportions of genes supporting each of two alternative hypotheses in the Spatafora2016_46taxa_192genes and Subset_Dikaya data matrices. The GLS values for each gene in each data matrix are provided in Supplementary Table 5. We considered a gene with an absolute value of log-likelihood difference of two as a gene with strong ($|\Delta \ln L| > 2$) or weak ($|\Delta \ln L| < 2$) phylogenetic signal.
